# Independent effects of testosterone, estradiol, and sex chromosomes on gene expression in immune cells of trans- and cisgender individuals

**DOI:** 10.1101/2024.10.08.617275

**Authors:** Rebecca M. Harris, Troy Whitfield, Laura V. Blanton, Helen Skaletsky, Kai Blumen, Phoebe Hyland, Em McDermott, Kiana Summers, Jennifer F. Hughes, Emily Jackson, Petra Teglas, Bingrun Liu, Yee-Ming Chan, David C. Page

**Affiliations:** Division of Endocrinology, Boston Children’s Hospital, Boston, MA 02115, USA; Harvard Medical School, Boston, MA 02115, USA; Whitehead Institute, Cambridge, MA 02142, USA; Howard Hughes Medical Institute, Whitehead Institute, Cambridge, MA 02142, USA; Department of Biology, Massachusetts Institute of Technology, Cambridge, MA 02139, USA

**Keywords:** Sex hormones, testosterone, estradiol, sex chromosomes, X chromosome, sex differences, transgender, gender, peripheral blood mononuclear cell, single-cell RNA-sequencing

## Abstract

The origins of sex differences in human disease are elusive, in part because of difficulties in separating the effects of sex hormones and sex chromosomes. To separate these variables, we examined gene expression in four groups of trans- or cisgender individuals: XX individuals treated with exogenous testosterone (n=21), XY treated with exogenous estradiol (n=13), untreated XX (n=20), and untreated XY (n=15). We performed single-cell RNA-sequencing of 358,426 peripheral blood mononuclear cells. Across the autosomes, 8 genes responded with a significant change in expression to testosterone, 34 to estradiol, and 32 to sex chromosome complement with no overlap between the groups. No sex-chromosomal genes responded significantly to testosterone or estradiol, but X-linked genes responded to sex chromosome complement in a remarkably stable manner across cell types. Through leveraging a four-state study design, we successfully separated the independent actions of testosterone, estradiol, and sex chromosome complement on genome-wide gene expression in humans.

## Introduction

Biological sex influences healthy traits and diseases in every organ system and across all ages.^1–7^ These sex-differential phenotypes are commonly attributed to the effects of the sex hormones, testosterone and estradiol, and the sex chromosomes, X and Y.

Testosterone and estradiol were initially investigated in Western medicine in the 18^th^ and 19^th^ centuries through a series of experiments delving into their paracrine effects through the removal^8,9^ and reimplantation of the gonads,^9–13^ and the use of gonadal extracts.^14–16^ As these hormones are synthesized in the gonads, it is not surprising that their identification and initial investigations into their actions were viewed through a reproductive-centric lens focused on the management of hot flashes, ovulation, and dysmenorrhea in women and listlessness in men.^17–19^ More recently, the roles of testosterone and estradiol in non-reproductive tissues have been studied more extensively, yielding insights into their effects in a wide range of tissues throughout the body, including but not limited to the brain,^20^ heart,^21–23^ vasculature,^21^ muscle,^24^ and adipose.^25^ The sex chromosomes originated as ordinary autosomes approximately 180 million years ago.^26,27^ Since that time, they have evolved into the differentiated sex chromosomes known today, the largest source of variation in the human genome. Females and males each have an equivalent “active” X chromosome (Xa) but differ in their second sex chromosome: females have an “inactive” X (Xi) and males have a Y. Similar to the sex hormones, historically, the effects of the sex chromosomes were also thought to be limited to the reproductive tract, with little impact in non-reproductive organs. This reproductive-centric view stemmed from two findings: 1) the second X chromosome in 46,XX cells underwent condensation and transcriptional attenuation due to X chromosome inactivation (XCI)^28,29^ and 2) Y chromosome genes are comparatively few in number and mainly expressed in the testes.^30,31^ However, multiple studies have shown that a significant fraction of genes on the human Xi (up to 32%) are expressed in both non-reproductive and reproductive tissues.^32–39^ On the Y chromosome, a subset of genes are broadly-expressed across tissues and are key regulators of transcription, translation, and protein stability.^40^ Xi and Y copy number affect the expression of 21% of autosomal genes,^41^ implying that broadly expressed genes encoded by Xi and Y have a widespread impact across the genome.^39^

Testosterone, estradiol, and the sex chromosomes exert their effects through changes in gene expression,^42–45^ therefore sex-differential gene expression is a molecular intermediary connecting the biological actions of testosterone, estradiol, and the sex chromosomes to their phenotypic effects. A major challenge in the field of sex-differential biology is how to distinguish the precise effects of testosterone, estradiol, and the sex chromosomes as they are typically linked: in placental mammals, the presence of an intact Y chromosome leads to the development of testes, which produce testosterone, while the absence of a Y chromosome leads to the development of ovaries, which produce estradiol.

To isolate the effects of testosterone, estradiol, and sex chromosome complement (XX or XY) in humans, we implemented a four-state, cross-sectional study design of trans- and cisgender individuals, the former receiving exogenous testosterone or estradiol for at least a year as part of their gender-affirming medical care. This study design allowed us to compare four combinations of sex hormones and sex chromosomes in humans: XX individuals treated with exogenous testosterone (henceforth “XX+T”), XY individuals treated with exogenous estradiol (“XY+E”), untreated XX individuals (“XX”), and untreated XY individuals (“XY”).

We chose to focus on the human immune system because it displays an impressively broad range of sex-differential biology,^46^ such as proportional abundances of immune cell types, responses to vaccination^47–51^ and infection,^52–57^ and rates of autoimmunity.^58^ We selected peripheral blood mononuclear cells (PBMCs) as our tissue of interest because PBMCs provide an easily accessible window into the immune system and have also been used as a microcosm that reflects changes in less accessible organ systems.^59–61^ Moreover, PBMCs are primary cells and therefore recapitulate *in vivo* physiology,^62^ exhibit sex-differential gene expression,^63^ and express the androgen receptor and both forms of the estrogen receptors.^64,65^

We performed single-cell RNA-sequencing (scRNA-seq) on 358,426 PBMCs isolated from 34 transgender and 35 cisgender individuals and identified 18 distinct cell types. We found that testosterone, estradiol, and sex chromosome complement have unique, non-overlapping effects on PBMC cell abundances and gene expression. Testosterone and estradiol influence abundances of PBMC cell types but sex chromosome complement does not. Within each cell type, we quantified gene expression at the levels of individual genes and functional gene-set enrichment, and identified non-overlapping sets of autosomal genes that significantly respond to a change in testosterone, estradiol, or sex chromosome complement. We also determined that the interferon alpha (IFNɑ) and gamma (IFNƔ) response pathways, which play key roles in autoimmunity and inflammation, responded positively to increasing testosterone concentration and negatively to increasing estradiol concentration. The IFNɑ and IFNƔ response pathways also responded positively to XY sex chromosome complement compared to XX. Across all 18 cell types, we determined that the autosomal gene responses to testosterone, estradiol, and sex chromosome complement are cell-type-specific, in contrast to the responses of X-chromosomal genes to sex chromosome complement, which are preserved across cell types. In fact, we found that genes expressed from Xi and also expressed in a wide array of cells have particularly stable responses to sex chromosome complement. Interestingly, we did not identify any sex chromosomal genes that responded to testosterone or estradiol.

Our study answers basic biological questions about the independent effects of testosterone, estradiol, and sex chromosome complement on immune cells, providing new mechanistic insights into the basis of the sex-differential nature of the human immune system.

## Results

### Human cohort with four combinations of sex hormones and sex chromosomes

To dissect out the independent effects of testosterone, estradiol, and sex chromosome complement on gene expression, we enrolled 69 individuals representing four combinations of sex hormones and sex chromosomes: 21 XX+T, 13 XY+E, 20 XX, and 15 XY (Figure 1). As part of the study, each individual provided self-reported demographic data, and medical information was extracted from the electronic medical record (Table 1).

**Figure 1.**
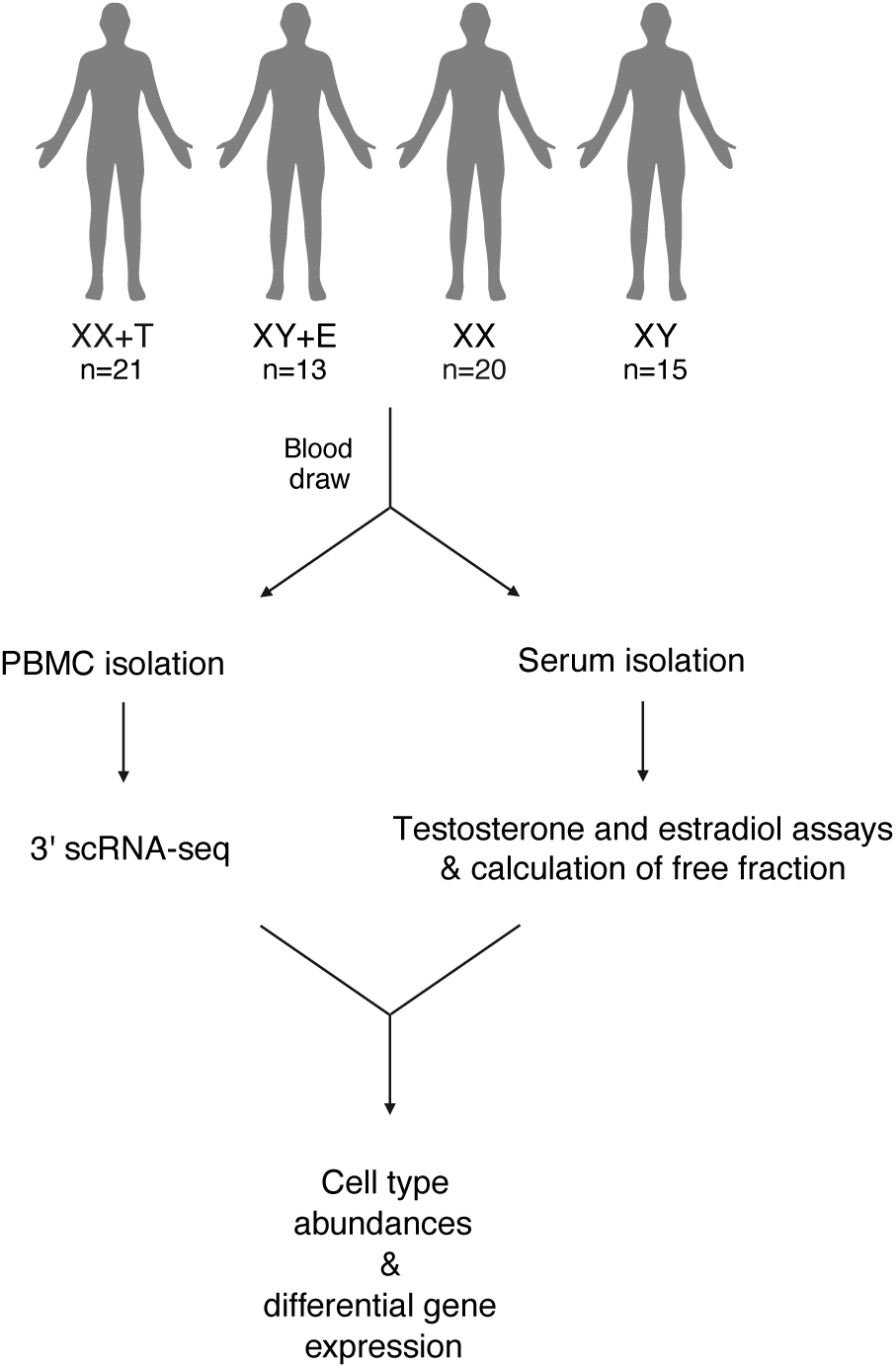
Human cohort with four combinations of sex hormones and sex chromosomes. In our study we included 34 transgender individuals receiving either exogenous testosterone (T) or estradiol (E) as part of their gender-affirming medical treatment for at least one year (XX+T and XY+E) and 35 cisgender individuals (XX and XY). We investigated the effects of testosterone, estradiol, and sex chromosome complement on peripheral blood mononuclear cell (PBMC) abundances and, within each cell type, differential gene expression at the single-cell level. See also Figure S1.

**Table 1.** Cohort description including 34 trans- and 35 cisgender individuals.

|  | XX+T | XY+E | XX | XY |
| --- | --- | --- | --- | --- |
| Number of individuals | 21 | 13 | 20 | 15 |
| Age (years) | 19.7 (2.2) | 19.2 (2.2) | 21.6 (3.8) | 23.1 (3.5) |
| BMI (kg/m<sup>2</sup>) | 26.9 (7.9) | 27.7 (12.5) | 22.9 (3.9) | 23.7 (1.8) |
| Race (self-reported) |  |  |  |  |
| Asian | 0 | 0 | 5 (25%) | 4 (27%) |
| Black/African American | 0 | 0 | 2 (10%) | 0 |
| White/Caucasian | 19 (90%) | 13 (100%) | 12 (60%) | 11 (73%) |
| Multiracial | 2 (10%) | 0 | 1 (5%) | 0 |
| Ethnicity (self-reported) |  |  |  |  |
| Hispanic/Latino/a/x/e | 4 (19%) | 0 | 3 (15%) | 2 (13%) |
| Not Hispanic/Latino/a/x | 17 (81%) | 13 (100%) | 17 (85%) | 13 (87%) |
| Gender (self-reported) |  |  |  |  |
| Female | 0 | 8 | 20 | 0 |
| Male | 10 | 0 | 0 | 15 |
| Non-binary/gender queer/gender fluid | 1 | 0 | 0 | 0 |
| Transgender female | 0 | 10 | 0 | 0 |
| Transgender male | 17 | 0 | 0 | 0 |
| GnRHa treatment prior to sample |  |  |  |  |
| None | 15 (100%) | 7 (54%) | - | - |
| Initiated in early puberty | 0 | 1 (8%) | - | - |
| Initiated in late puberty | 0 | 5 (38%) | - | - |
| Gonadectomy prior to sample | 0 | 2 (15%) | - | - |
| Initiation of exogenous sex hormone treatment |  |  |  |  |
| Early puberty | 0 | 1 (8%) | - | - |
| Late puberty | 21 (100%) | 12 (92%) | - | - |
| Duration of exogenous sex hormone treatment (years) | 3.1 (1.7) | 2.5 (1.5) | - | - |
| Additional hormonal medications at time of sample |  |  |  |  |
| None | 15 (100%) | 2 (15%) | 20 (100%) | 15 (100%) |
| GnRHa | - | 4 (31%) | - | - |
| Spironolactone | - | 7 (54%) | - | - |

Age, body mass index (BMI), and medical treatment were obtained from review of the electronic
medical record. Race, ethnicity, and gender were self-reported via questionnaire. Individuals
were allowed to select more than one gender from a list of options and were also allowed to
write in their own response. Absolute numbers and percentages are noted; for age and BMI the
standard deviation is noted in the parentheses. Abbreviations: testosterone (T), estradiol (E),
and gonadotropin releasing hormone agonist (GnRHa).

We collected blood samples from each individual and isolated PBMCs and serum the same day. We profiled PBMC gene expression in each individual using the 10x Genomics 3’ scRNA-seq Platform and generated 150-bp paired-end reads with an average of 48,007 reads/cell. From serum, we measured levels of total testosterone (Figure S1A), total estradiol (Figure S1B), and sex hormone binding globulin. We then calculated the “free” fraction of testosterone and estradiol to better reflect the concentration of bioavailable/active sex hormone (Figure S1C-D).^66^ The total and free levels of each sex hormone were highly correlated (Figure S1E-F).

### An inclusive model to dissect the independent effects of testosterone, estradiol, and sex chromosome complement

To maximize statistical power, we analyzed all 69 individuals together, in one model, because such an inclusive model provides more power than pairwise comparisons.^39,41^ This is particularly important when studying the effects of biological sex on gene expression, as sex differences in gene transcription are subtle.^39,41,43^ We tested the independent effects of testosterone, estradiol, and sex chromosome complement on PBMC cell type abundances using a linear mixed effects model. Then, within each cell type, we tested the effects of the same covariates on gene expression using the generalized linear model framework of DESeq2. We modeled calculated free testosterone and estradiol as continuous covariates and sex chromosome complement as a categorical variable. We standardized testosterone, estradiol, and sex chromosome complement to allow for comparison of responses. We also included age^67–69^ and BMI^70–72^ as continuous covariates, as each has known effects on PBMC gene expression, as well as emulsion batch as a categorical covariate.

We sequenced RNA from a total of 358,426 PBMCs, with an average of 5,195 ± 1,741 cells/individual. We identified 18 cell types (Figure 2A) using known cell type markers (Table S1) and, for rarer cell types, via alignment to a single-cell PBMC reference atlas.^73^

**Figure 2.**
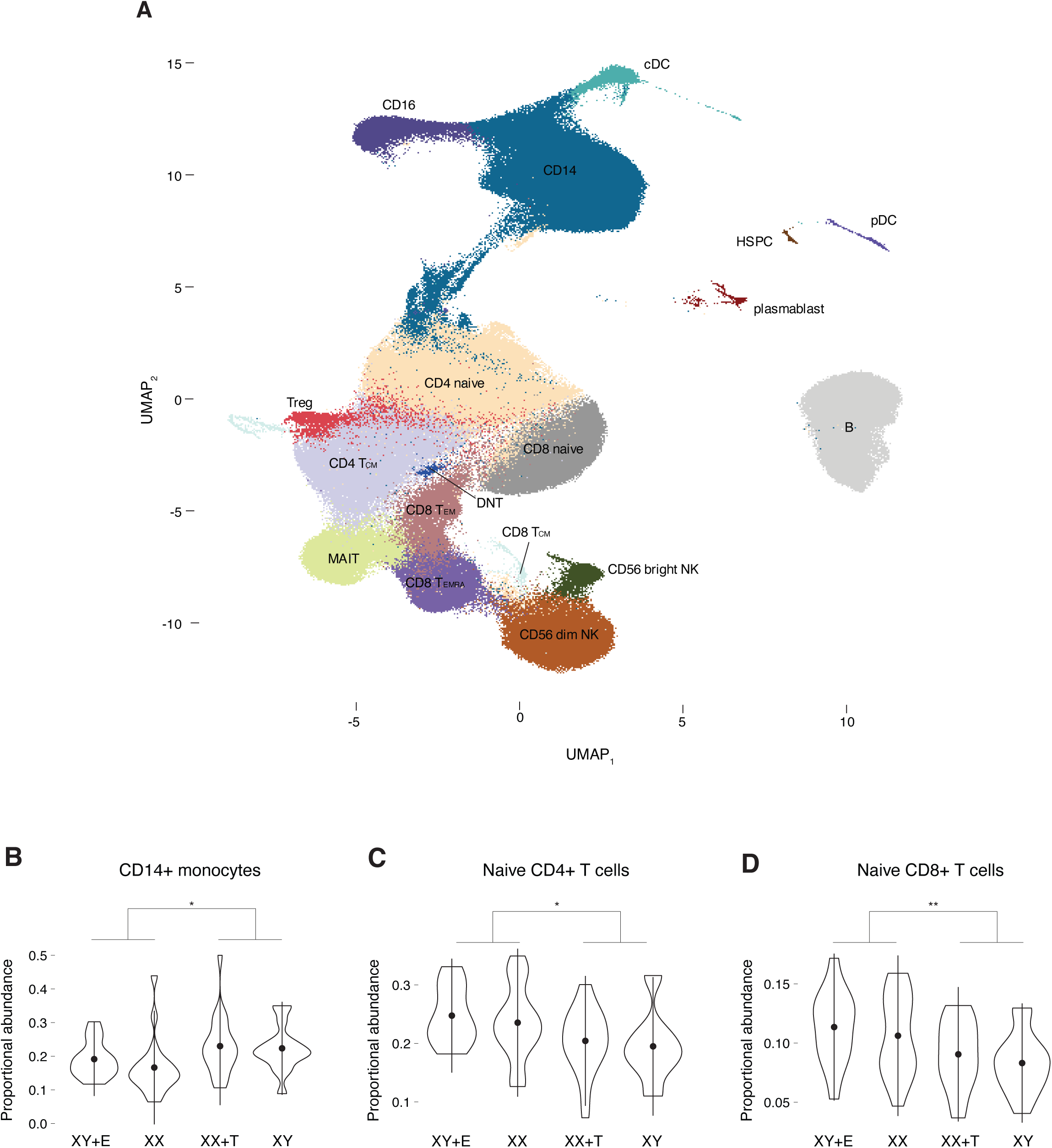
Testosterone influences abundances of CD14+ monocytes and naïve CD4+ and CD8+ T cells. **(A)** Uniform manifold approximation projection (UMAP) showing 18 distinct PBMC cell types. **(B-D)** Violin plots showing median (dot) and interquartile range (vertical lines) of the proportional abundances of CD14+ monocytes **(B)**, naïve CD4+ T cells **(C)**, and naïve CD8+ T cells **(D)** separated by group. Significant differences noted by asterisks (* *p*-adj <0.05; ** *p*-adj <0.01). See also Table S1 and Figure S2.

### Testosterone influences abundances of CD14+ monocytes and naïve CD4+ and CD8+ T cells

Sex differences in abundances of PBMC cell types occur throughout the lifespan.^74^ Our goal was to understand the basis for these sex differences by untangling the effects of testosterone, estradiol, and sex chromosome complement on changes in cell type abundance across the 18 cell types identified.

Importantly, we detected no significant effect of sex chromosome complement on the proportional abundance of any of the 18 cell types, but there were significantly different abundances of CD14+ monocytes, naïve CD4+ T cells, and naïve CD8+ T cells when comparing estradiol-predominant (XY+E and XX) and testosterone-predominant (XX+T and XY) individuals. We found that the average abundance of CD14+ monocytes was 29% higher in the testosterone-predominant groups compared with the estradiol-predominant groups (p<0.031) (Figure 2B). Both testosterone and estradiol were associated with an increase in CD14+ monocyte cell abundance (testosterone p=0.004, estradiol p=0.03) (Figure S2A,D), but testosterone explained more of the variance than estradiol (testosterone R^2^=0.12, estradiol R^2^=0.07) (Figure S2A,D).

Testosterone was the only modulator of naïve CD4+ and CD8+ T cell abundances. In naïve CD4+ T cells, the testosterone-predominant groups had, on average, a 16.5% decrease abundance relative to the estradiol-predominant groups (p<0.013) (Figure 2C). While testosterone concentration significantly correlated with naïve CD4+ T cell abundance, explaining 8% of the variance (R^2^=0.08, p=0.02), abundance did not vary significantly with estradiol concentration, suggesting that testosterone, but not estradiol, modulates naïve CD4+ T cell abundance (Figure S2B,E). We then completed the same analysis in naïve CD8+ T cells and found that the average cell abundance was 19.8% lower in the testosterone-predominant groups compared with the estradiol-predominant groups (p<0.0064) (Figure 2D). Testosterone explained 11% of the variance in naïve CD8+ T cell abundance (R^2^=0.11, p=0.006), while abundance did not vary significantly with estradiol concentration (Figure S2C,F).

Overall, testosterone had modest but significant effects on abundances of CD14+ monocytes and naïve CD4+ and CD8+ T cells. Our findings align well with observations in other clinical settings: One study of healthy men undergoing androgen-blockade with a gonadotropin releasing hormone agonist (GnRHa) followed by testosterone replacement found that testosterone increased monocyte abundance,^75^ although the study did not distinguish between CD14+ and CD16+ monocytes. Another study demonstrated that two men with hypogonadism had increased abundances of naïve CD4+ T cells; one of the men underwent treatment with testosterone and the abundance of naïve CD4+ T cells decreased.^76^ Finally, a study of sixteen men with prostate cancer found that naïve CD4+ and CD8+ T cell abundances increased after androgen-blockade with a GnRHa.^77^ Our study builds upon these findings and confirms that it is testosterone, independent of sex chromosome complement, that impacts cell abundance, exerting effects on both the innate and adaptive arms of the immune system.

### Autosomal responses to testosterone, estradiol, and sex chromosome complement are cell-type-specific

Our lab previously demonstrated that X and Y chromosome dosage are associated with distinct autosomal responses in four different cell types, including two types of primary immune cells (CD4+ T cells and monocytes).^78^ We wanted to expand upon this finding and investigate whether autosomal responses to testosterone, estradiol, and sex chromosome complement are also cell-type-specific using the 18 cell types in our dataset.

Given that X- and Y-chromosomal gene expression differs between XX and XY individuals due to differences in X and Y chromosome counts, yet all individuals possess the same number of autosomes, we examined autosomal and sex-chromosomal gene responses separately. Across the 18 cell types, we identified a total of 74 autosomal genes with significant responses (i.e. log_2_ fold change significantly different from zero) to either testosterone, estradiol, or sex chromosome complement (Figures 3A-C, Tables S2-S3). Within each cell type, we did not identify any autosomal genes with a significant response to more than one covariate.

**Figure 3.**
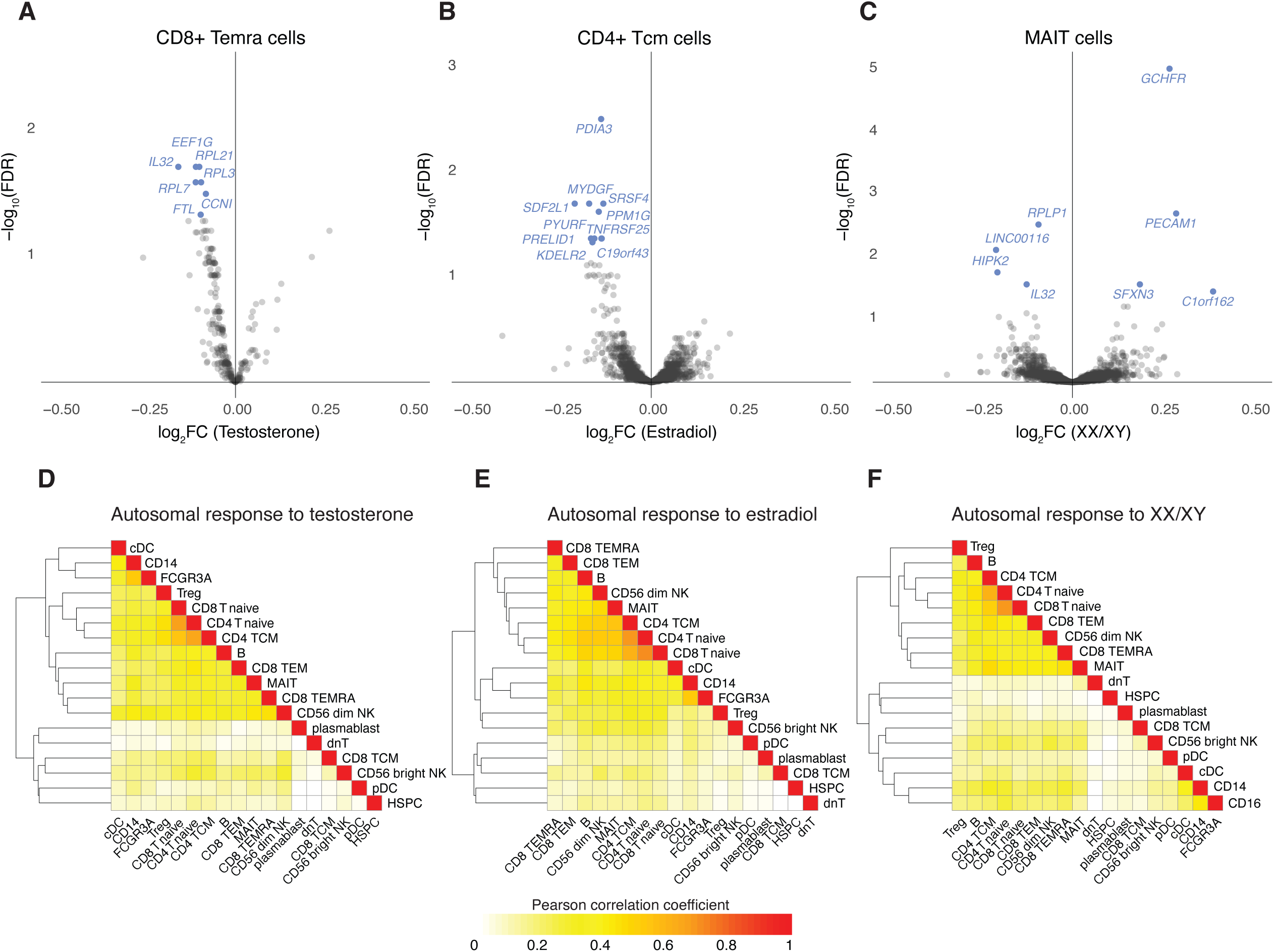
Autosomal responses to testosterone, estradiol, and sex chromosome complement are cell-type-specific. **(A-C)** Volcano plots of the autosomal responses (log_2_ fold change (log_2_FC) in expression) to testosterone in CD8+ Temra cells **(A)**, estradiol in CD4+ Tcm cells **(B)**, and sex chromosome complement (XX/XY) in MAIT cells **(C)**. Genes with a significant response (FDR <0.05) are noted in blue. Heatmaps of Pearson correlation coefficients of the autosomal responses to testosterone **(D)**, estradiol **(E)**, and sex chromosome complement **(F)** between every pair of cell types. See also Tables S2-S4.

Eight autosomal genes across two cell types had significant responses (FDR <0.05) to testosterone (Figure 3A, Table S2). Seven genes had a negative response with increasing testosterone concentration and one gene had a positive response. Thirty-four autosomal genes across six cell types had significant responses to estradiol (Figure 3B, Table S2). Thirty-one out of the 34 genes had a negative response with increasing estradiol concentration and three genes had a positive response. Twenty-seven of the 34 genes (79%) had a response in only one cell type, highlighting the diversity of the autosomal transcriptome between different cell types. Interestingly, the number of autosomal genes with a significant response to estradiol was four times the number with a significant response to testosterone.

Thirty-two autosomal genes across 10 cell types had a significant response to sex chromosome complement (Figure 3C, Table S2). Twenty-nine of the 32 genes (91%) had a significant response in only one cell type, demonstrating that the autosomal response to sex chromosome complement across cell types exhibited the same degree of diversity as the testosterone and estradiol responses. Out of the genes that had a significant response in more than one cell type, sideroflexin 3 (*SFXN3*) was particularly interesting because its expression increased in XX individuals compared to XY individuals in the greatest number of cell types (6/18) (Table S2). *SFXN3* encodes a serine transporter located on the inner mitochondrial membrane, which is involved in one-carbon metabolism and the production of glycine.^79^ In cancer, products of one-carbon metabolism are substrates for tumor cell proliferation.^80–82^ Sex differences in one-carbon metabolism are typically attributed to the sex hormones.^83–85^ Our results suggest a potential role for sex chromosome complement, independent of sex hormones, in the one-carbon metabolism pathway.

To confirm that the autosomal responses to testosterone, estradiol, and sex chromosome complement were cell-type-specific, we calculated the Pearson correlation coefficients of the responses to testosterone (Figure 3D), estradiol (Figure 3E), and sex chromosome complement (Figure 3F) for all autosomal genes expressed in each pair of cell types. We found that the Pearson correlation coefficients for the response of autosomal genes to testosterone, estradiol, and sex chromosome complement were low (testosterone 0.18±0.14, estradiol 0.2±0.15, sex chromosome complement 0.15±0.14 (Figure 5F), which demonstrates the cell-type-specific nature of the autosomal responses.

We also verified that the cell-type-specific nature of the autosomal responses to testosterone, estradiol, and sex chromosome complement was not due to a lack of overlap in expressed autosomal genes across the 18 cell types (Tables S3-S4).

### Interferon response pathways are altered by testosterone, estradiol, and sex chromosome complement

As responses of individual genes to biological sex are often subtle,^43^ we chose to take the powerful approach of investigating the responses of entire gene sets to elucidate the roles of testosterone, estradiol, and sex chromosome complement in key biological processes. Within each cell type, we used gene set enrichment analysis (GSEA)^86^ of the Hallmark collection of the Molecular Signatures Database^87^ to identify gene sets enriched in response to testosterone, estradiol, or sex chromosome complement (Figures 4A-C). These highly curated gene sets represent a wide array of key biological pathways. As autosomal and sex-chromosomal gene responses to biological sex manifest differently, we removed the sex-chromosomal genes from each gene set to focus on autosomal genes. We carried out GSEA on 900 cell-type:gene-set pairs (18 cell types x 50 gene sets). We determined that the combined effects of testosterone, estradiol, and sex chromosome complement are associated with significant alterations in the enrichment of 64% (32/50) of the pathways. Additionally, we found that testosterone and estradiol are associated with enrichment of a greater number of pathways than sex chromosome complement (testosterone 50% (25/50); estradiol 46% (23/50); sex chromosome complement 22% (11/50)).

**Figure 4.**
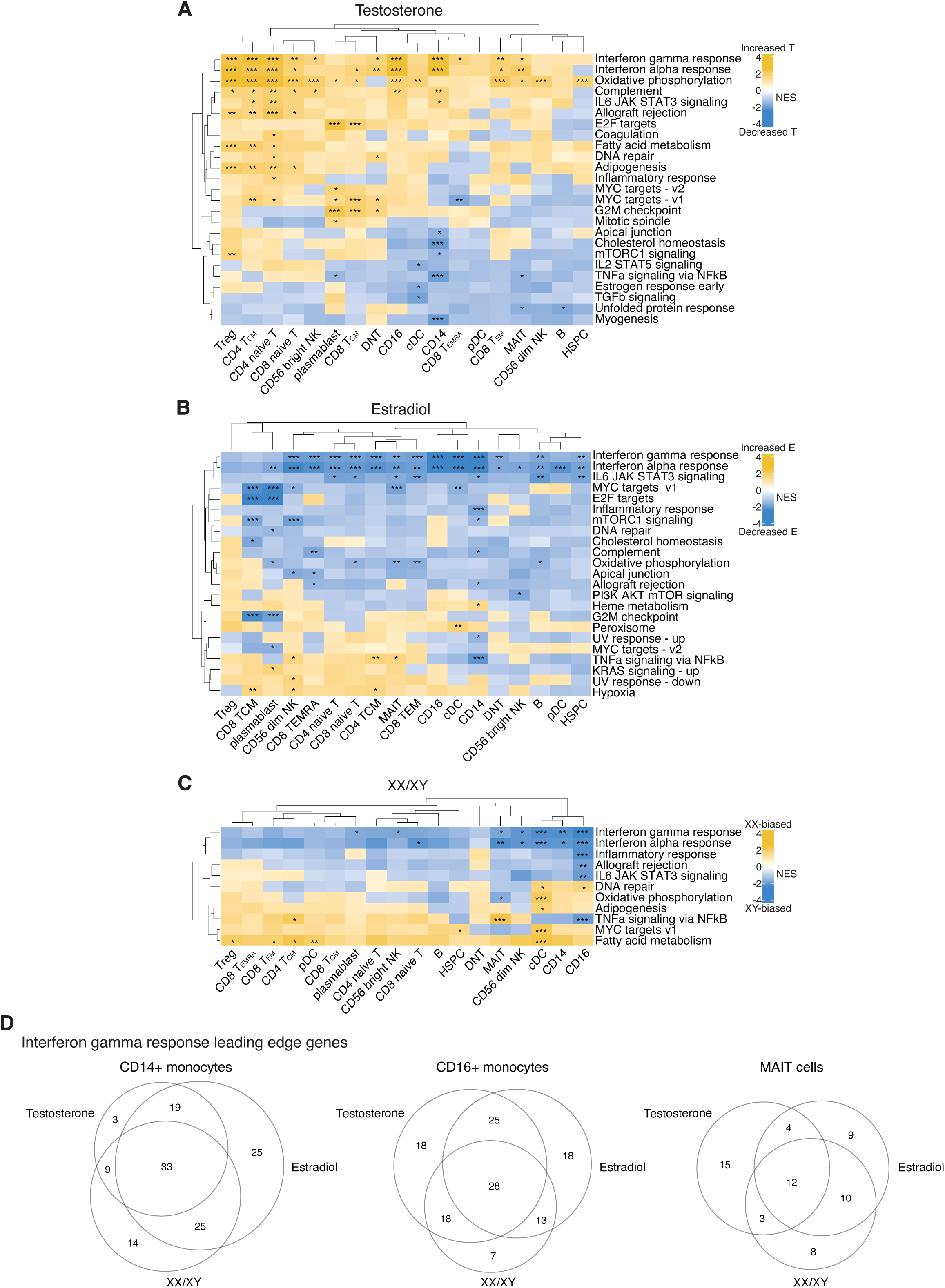
Interferon response pathways are altered by testosterone, estradiol, and sex chromosome complement. **(A-C)** Heatmaps of normalized enrichment scores (NES) for Hallmark gene sets in all 18 cell types based on the effects of testosterone **(A)**, estradiol **(B)**, and sex chromosome complement **(C)**. Significant enrichments noted by asterisks (* *p*-adj <0.05; ** *p*-adj <0.01; *** *p*-adj <0.001). **(D)** Venn diagrams of the number of interferon gamma response leading edge genes specific to testosterone, estradiol, or sex chromosome complement, or shared, in CD14+ monocytes (left), CD16+ monocytes (middle), and MAIT cells (right).

Across all fifty pathways, the interferon alpha (IFNɑ) and gamma (IFNƔ) response pathways stand out as the pathways with significant responses to testosterone, estradiol, and sex chromosome complement across the greatest number of cell types (Figure 4A-C). We found that 56% (10/18) of cell types had significantly positive enrichments in the IFNɑ pathway in response to increasing testosterone concentration and 61% (11/18) of cell types had significantly positive enrichments in the IFNƔ pathway (Figure 4A). The opposite response occurred with estradiol – 89% (16/18) of cell types had significantly negative enrichments in the IFNɑ pathway in response to increasing estradiol concentration and 72% (13/18) of cell types had significantly negative enrichments in the IFNƔ pathway (Figure 4B). We also found that the IFNɑ and IFNƔ pathways had significantly positive enrichments with XY sex chromosome complement compared to XX in 33% (6/18) and 39% (7/18) of cell types, respectively (Figure 4C).

Since the IFNɑ and IFNƔ response pathways had significantly positive enrichments with increasing testosterone concentration and also with XY sex chromosome complement, we wondered whether the leading-edge genes, which are the genes within a given pathway that drive the enrichment, were the same or different between testosterone, estradiol, and sex chromosome complement. The IFNƔ pathway was significantly enriched by all three covariates in CD14+ monocytes, CD16+ monocytes, and MAIT cells, so we focused our analysis on those three cell types. Across the three cell types, we found that the majority of leading-edge genes responded to only one covariate (33-52% responded to one covariate; 28-44% responded to two covariates; and 20-26% responded to three covariates) (Figure 4D).

The IFN pathways have known sex-biases: peripheral blood lymphocytes from healthy women produce more IFNɑ upon stimulation with Toll-like receptor 7 than peripheral blood lymphocytes from healthy men^88^ and naïve CD4+ T cells from healthy women produce more IFNƔ in response to anti-CD3 and anti-CD28 than naïve CD4+ T cells from healthy men.^89^ Increased production of IFNɑ^90^ and IFNƔ^91^ cytokines and positive enrichment of the IFNɑ^90,92^ and IFNƔ^93^ response pathways are associated with the pathogenesis of autoimmune diseases. Autoimmune diseases demonstrate a particularly high degree of female sex-bias,^58^ and increased production of IFNɑ^90^ and IFNƔ^91^ in women with autoimmune disease has been implicated as a potential cause of the female sex-bias. Despite all of this evidence, the distinct roles of testosterone, estradiol, and sex chromosome complement in the healthy, unstimulated state, and in autoimmune diseases have been challenging to disentangle. Here, we are able to separate out the effects of testosterone, estradiol, and sex chromosome complement on the IFNɑ and IFNƔ response pathways in the healthy-state, providing new insights into baseline sex differences in the immune system, and potentially contributing to our knowledge about sex differences in autoimmune disease in the future.

### Responses of X-chromosomal genes to sex chromosome complement are preserved across 18 cell types

Our lab previously showed that X-chromosomal gene responses to Xi dosage are preserved across four cell types.^78^ We wanted to build upon this finding and investigate whether the responses of X-chromosomal genes to sex chromosome complement are more consistent than autosomal responses across our 18 cell types. Additionally, we wanted to determine whether X-chromosomal genes respond to testosterone or estradiol.

We specifically focused on genes located in the non-pseudoautosomal region of the X chromosome (NPX), a region that differs in gene content from the Y chromosome. We did not identify any NPX genes that responded to testosterone or estradiol. The androgen receptor (*AR*) is located on the X-chromosome and there are reports of downregulation of *AR* mRNA upon exposure to androgens in human prostate and breast cancer cell lines, upregulation in human osteosarcoma and sarcoma cell lines, and no effect in human skin fibroblasts.^94–96^ These previous studies suggest that androgen regulation of *AR* is variable and cell-type-specific, and our data showing a lack of response in PBMCs reflects this.

Across the 18 cell types, we identified 28 NPX genes that responded to sex chromosome complement (XX/XY) (Table S5-S6). Twenty-five of the 28 genes have previously been shown to be expressed from Xi,^39^ and as such have higher expression levels in XX compared to XY cells. As examples, 7% (19/290) of NPX genes expressed in CD56 dim NK cells and 6% (16/286) of NPX genes expressed in MAIT cells had increased expression in XX individuals compared to XY individuals (“XX-biased”) (Figures 5A,B; *XIST* not shown).

Overall, 27 out of the 28 significant NPX genes were XX-biased (Table S5). One NPX gene, *SCML1*, had higher expression in classical dendritic cells from XY individuals compared to XX individuals (“XY-biased”) (Table S5). This finding was consistent with previous *in vitro* studies using LCLs and fibroblasts, which found that expression of *SCML1* responds negatively to increasing X chromosome dosage.^41^

To clearly visualize the effects of different combinations of testosterone, estradiol, and sex chromosome complement on gene expression, we used our four-state study design to compare expression of broadly-expressed NPX genes across the four groups. We focused on two prominent NPX genes, *KDM6A* and *EIF2S3,* which had robust significant responses to sex chromosome complement across the majority of cell types (*KDM6A* 12/18 cell types; *EIF2S3* 16/18 cell types). We found that *KDM6A* and *EIF2S3* had increased expression in MAIT cells from XX individuals, regardless of sex hormonal milieu (*KDM6A* p=5.5e-17; *EIF2S3* p=6.2e-21) (Figure 5C). This finding is consistent with previous studies demonstrating that expression of *KDM6A* and *EIF2S3* responds in a dose-dependent manner to the number of X chromosomes.^39,78^

Many XX-biased NPX genes are highly evolutionarily conserved between species and are involved in key cellular processes (e.g. transcription, splicing, translation, chromatin modification, ubiquitination).^40^ Therefore, we hypothesized that for each NPX gene, the response to sex chromosome complement would be similar across cell types. We found that there was a range of responses of NPX genes to sex chromosome complement across cell types (Figure 5D; *XIST* not shown). We calculated the Pearson correlation coefficient of the response of expressed NPX genes to sex chromosome complement for every pair of cell types (Figure 5E). We found that the correlation coefficients of the responses of NPX genes to sex chromosome complement were significantly greater than the correlation coefficients of autosomal responses to sex chromosome complement (Figure 5F). The correlation coefficients of the responses of NPX genes to sex chromosome complement were also significantly greater than those of the responses of NPX or autosomal genes to testosterone or estradiol (Figure 5F). These findings highlight that the responses of X-chromosomal genes to sex chromosome complement are preserved across cell types. We postulate that this stability is likely due to the key biological processes in which many NPX genes are involved.^40^

**Figure 5.**
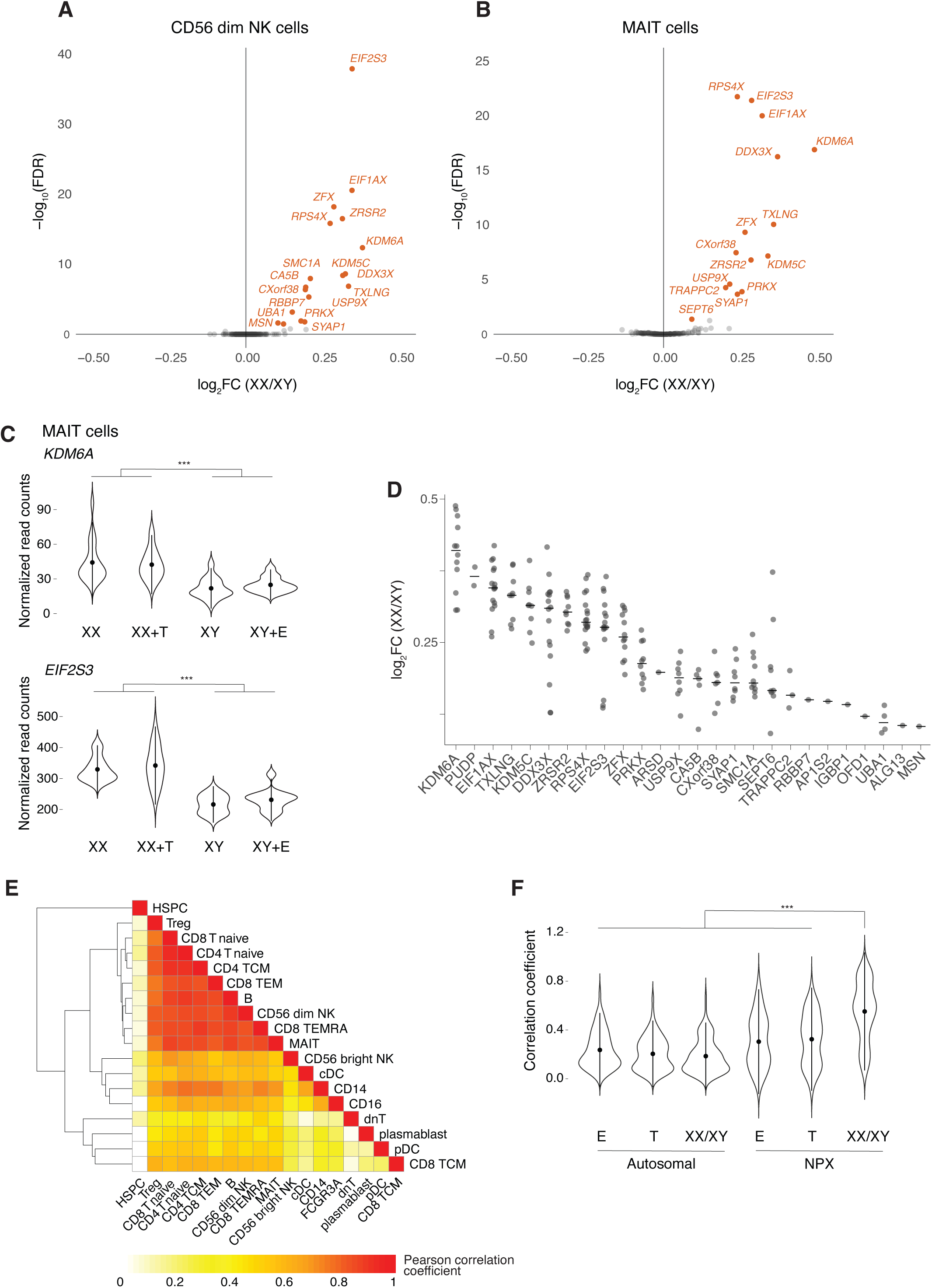
Responses of X-chromosomal genes to sex chromosome complement are preserved across 18 cell types. **(A-B)** Volcano plots of the responses of non-PAR X-chromosomal (NPX) genes to sex chromosome complement in CD56 dim natural killer (NK) **(A)** and mucosal associated invariant T (MAIT) **(B)** cells. Genes with a significant response (FDR <0.05) are noted in orange. **(C)** Violin plots showing the median (dot) and interquartile range (vertical lines) of normalized read counts of two representative NPX genes, *KDM6A* and *EIF2S3*, separated by group. FDR calculated from unpaired t-tests; significant differences noted by asterisks (*** p-adj <0.001). **(D)** Dot plot of the responses of significant NPX genes to sex chromosome complement. Each dot is the log_2_ fold change (log_2_FC) in expression of a significant gene (FDR<0.05) within a specific cell type; horizontal line is the median. **(E)** Heatmap of Pearson correlation coefficients of NPX responses to sex chromosome complement between every pair of cell types (excluded). **(F)** Violin plot showing the median (dot) and interquartile range (IQR; vertical lines) of the Pearson correlation coefficients comparing the responses of autosomal and NPX genes to testosterone, estradiol, and sex chromosome complement between every pair of cell types. Significant differences noted by asterisks (*** p-adj <0.001). See also Tables S5-S6.

### Responses of X-chromosomal genes to sex chromosome complement are more stable than responses of autosomal genes

To further dissect out the underlying etiology of our finding that responses of X-chromosomal genes to sex chromosome complement are preserved across different cell types compared to responses of autosomal genes, we utilized a quantitative metric: the coefficient of variation (CV).^78^ The CV is the ratio of the standard deviation to the mean, and in our analysis, the CV measures the degree of variation in the response of a given gene to sex chromosome complement across cell types. Therefore, a lower CV denotes less variability, or more stability, in gene response across cell types.

We first asked whether there was a relationship between the responses of genes to sex chromosome complement and the CV. Specifically, we hypothesized that genes with responses that differed the most between XX and XY individuals (i.e. genes with the greatest absolute effect sizes) would have more stable responses across cell types (i.e. lower CVs). We found that for both NPX and autosomal genes, genes with expression that differed the most between XX and XY individuals carefully maintained that difference in expression across cell types (Figure 6A).

**Figure 6.**
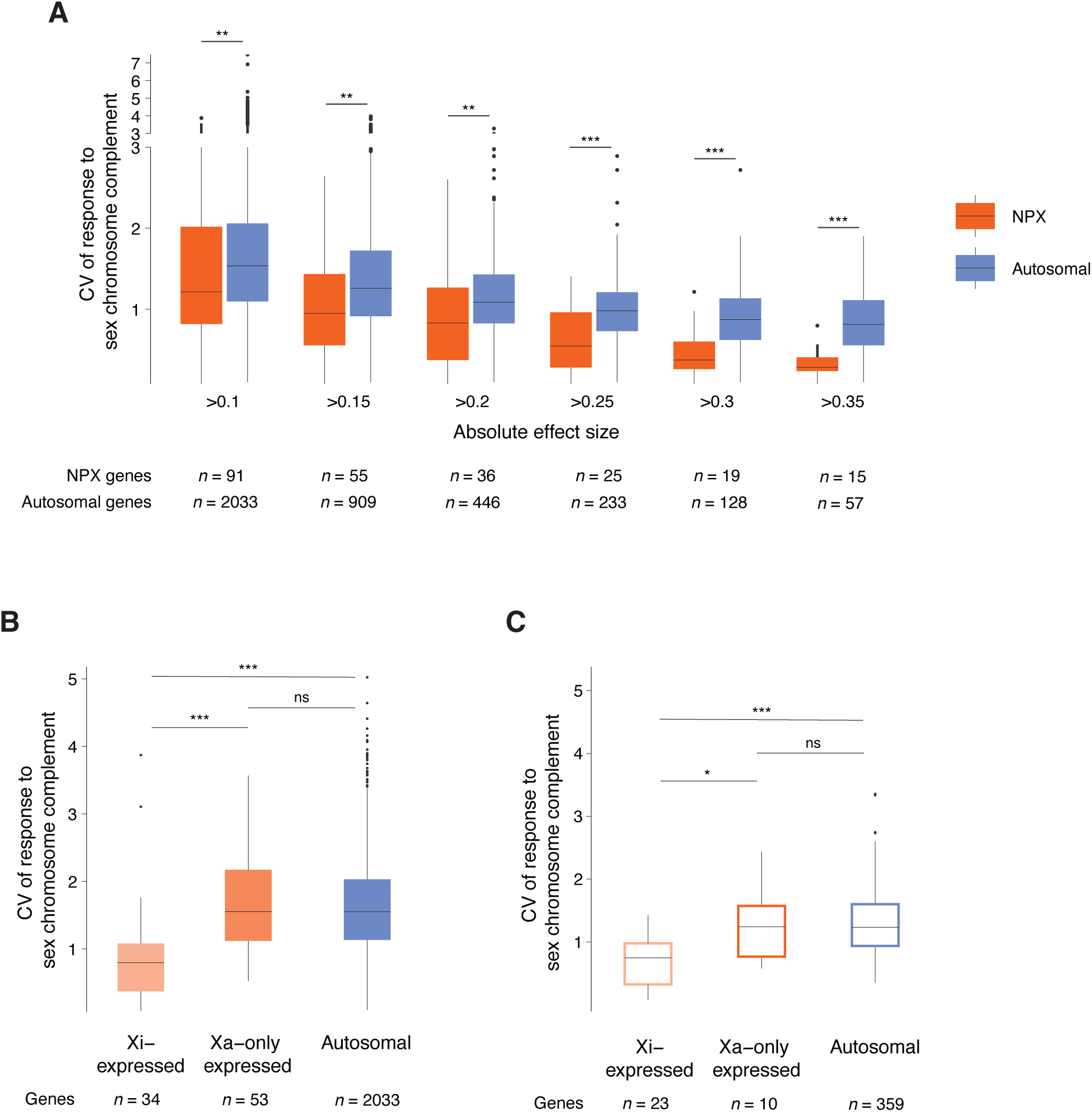
Responses of broadly expressed Xi-expressed genes to sex chromosome complement are preserved across cell types. **(A)** Variation in gene-by-gene responses to sex chromosome complement across all 18 cell types in which the gene was expressed. Variation was calculated using the coefficient of variation (CV = standard deviation in effect size across cell types / absolute mean effect size across cell types). Box plots show median (line), interquartile range (IQR; top and bottom of box), and 1.5 * IQR (whiskers) of CV values for non-pseudoautosomal region X-chromosomal (NPX) (orange) and autosomal (blue) genes. Gene sets restricted by absolute mean effect sizes above the indicated thresholds. FDR calculated from unpaired t-tests and asterisks indicate significance (** *p*-adj <0.01; *** p-adj <0.001). **(B)** Variation in responses to sex chromosome complement for genes expressed from the inactive and active X chromosomes (Xi-expressed; light orange), only the active X chromosome (Xa-only expressed; dark orange), or autosomes (blue). Analysis limited to genes expressed in at least 5 cell types. Box plots show median (line), interquartile range (IQR; top and bottom of box), and 1.5 * IQR (whiskers) of CV values. FDR calculated from unpaired t-tests and asterisks indicate significance (*** p-adj <0.001). **(C)** Variation in responses to sex chromosome complement for genes expressed from the indicated groups that are broadly expressed in 17 or 18 cell types. Box plots show median (line), interquartile range (IQR; top and bottom of box), and 1.5 * IQR (whiskers) of CV values. FDR calculated from unpaired t-tests and asterisks indicate significance (* *p*-adj <0.05; *** p-adj <0.001).

We also found that for a given absolute effect size, the responses of NPX genes to sex chromosome complement were more stable across cell types than responses of autosomal genes (Figure 6A). Additionally, the greater the absolute effect size, the greater the difference in the mean CV between NPX and autosomal genes. Meaning that for genes that differ the most in their responses between XX and XY participants, the NPX genes displayed significantly greater stability in response than autosomal genes. NPX genes with the greatest absolute effect sizes (>0.35) had 2.4 times more stable responses to sex chromosome complement than the responses of autosomal genes.

### Responses of Xi-expressed genes to sex chromosome complement are highly stable

We next investigated which NPX genes drive the particularly stable response to sex chromosome complement. First, we separated the NPX genes into two, non-overlapping categories: 1) genes expressed from Xi and Xa (“Xi-expressed”) and 2) genes expressed only from Xa but not Xi (“Xa-only expressed”).^39,78^ Based on our previous results studying the effects of Xi and Y dosage, we hypothesized that across our 18 cell types, Xi-expressed genes would have more stable responses to sex chromosome complement than Xa-only expressed or autosomal genes.

We calculated the CV for the responses of Xi-expressed, Xa-only expressed, and autosomal genes to sex chromosome complement and found that the responses of Xi-expressed genes have significantly lower CVs compared to Xa-only expressed and autosomal genes (Figure 6B). In fact, the responses of the Xa-only and autosomal genes did not have significantly different CVs (Figure 6B). This finding highlights the stability and cell-type-agnostic nature of the responses of Xi-expressed genes to sex chromosome complement while also demonstrating the cell-type-specific Xa-only and autosomal gene responses to sex chromosome complement.

### Responses of broadly expressed Xi-expressed genes to sex chromosome complement are preserved across cell types

We next asked whether genes that are more broadly expressed, in 17 or 18 PBMC cell types, have particularly stable responses to sex chromosome complement. Specifically, we hypothesized that broadly expressed Xi-expressed genes, which are dosage-sensitive and involved in key biological processes utilized by all cells (e.g. transcription, translation, epigenetic modification, etc.),^40^ may exhibit more precise maintenance of their responses to sex chromosome complement than broadly expressed Xa-only or autosomally expressed genes.

To answer this question, we calculated the CVs of the responses of Xi-expressed, Xa-only expressed, and autosomal genes that are broadly expressed in 17 or 18 cell types. We found that the broadly expressed Xi-expressed genes have significantly more stable responses to sex chromosome complement than broadly expressed Xa-only and autosomal genes (Figure 6C). Broadly expressed Xa-only genes did not have significantly different responses to sex chromosome complement compared to autosomal genes, highlighting that the responses of Xa-only and autosomal genes to sex chromosome complement are more cell-type-specific. These findings again distinguish the Xi-expressed genes and show that precise regulation of Xi-expressed gene responses to sex chromosome complement is preserved across cell types.

## Discussion

Our study included four combinations of sex hormones and sex chromosomes in humans, which allowed us to quantitatively distinguish the independent actions of testosterone, estradiol, and sex chromosome complement *in vivo* in 18 types of primary immune cells. We confirmed previous findings in the literature that testosterone increases monocyte^75^ and decreases naïve T cell^76,77^ proportional abundances and also further demonstrated that the effect of testosterone on PBMC abundance is independent of sex chromosome complement. We then used generalized linear modeling to characterize the effects of testosterone, estradiol, and sex chromosome complement on a gene-by-gene basis transcriptome-wide.

### Autosomal responses to testosterone, estradiol, and sex chromosome complement are distinct and cell-type-specific

Our four-state study design allowed us to delineate the independent effects of testosterone, estradiol, and sex chromosome complement on gene expression. We found that the autosomal genes with significant responses to testosterone, estradiol, and sex chromosome complement are distinct across the 18 cell types. We did not identify any autosomal genes with a significant response to a combination of two or three of the covariates. Interestingly, we also did not identify any sex-chromosomal genes that responded significantly to testosterone or estradiol.

### Responses of broadly expressed Xi-expressed genes to sex chromosome complement are preserved across cell types

We determined that the X-chromosomal genes with the greatest difference in expression between XX and XY individuals (absolute log_2_ fold change >0.35) are significantly more stable in their responses to sex chromosome complement than autosomal genes with the equivalent responses to sex chromosome complement. Delving deeper into genes expressed from Xi versus only expressed from Xa, we found that it is 23 broadly expressed Xi-expressed genes that have particularly stable responses to sex chromosome complement. Thirteen of the 23 broadly-expressed Xi-expressed genes have Y homologs in humans (*DDX3X, EIF1AX, KDM6A, PRKX, RPS4X, TXLNG, USP9X,* and *ZFX*) or other species (*AP1S2*, *EIF2S3X, P2RY8, UBA1,* and *ZRSR2*).^40,97^ These X-Y pair genes encode proteins that are necessary for key biological processes such as transcription, translation, and epigenetic modification.^40^ The ubiquitous need of all cells to carefully choreograph these key biological processes is likely the underlying reason for the increased stability in expression. The remaining ten genes are *ALG13, CA5B, CD99, CXorf38, NAA10, RBBP7, SMC1A, SYAP1, TRAPPC2,* and *XIST*. We postulate that these 23 genes, which are widely expressed across different tissues, may contribute to the phenotypic differences seen in XX and XY individuals throughout the body in health and disease.

## Conclusion

Through the implementation of an inclusive research design comprised of trans- and cisgender individuals, we successfully separated and identified the independent actions of testosterone, estradiol, and sex chromosome complement transcriptome-wide. A small number of translational studies have previously included transgender individuals, however, those studies mainly focused on investigating the longitudinal effects of exogenous testosterone in XX individuals or estradiol in XY individuals,^98–101^ not on dissecting the independent actions of testosterone, estradiol, and sex chromosome complement.

Elucidating the biological origins underlying sex differences in human health and disease is critical. As an example, it is well-established that autoimmune disease is strongly female-biased, however, it has been challenging to untangle the effects of testosterone, estradiol, and sex chromosome complement in autoimmunity. In the transgender population, there are reports of increased autoimmunity even before medical treatment^102–105^ and also in XY individuals after treatment with exogenous estradiol,^106–108^ therefore understanding the contributions of testosterone, estradiol, and sex chromosome complement to autoimmunity is vital. Using our four-state model, we determined that the IFNɑ and IFNƔ response pathways, which are implicated in autoimmunity, have positive responses to increasing testosterone concentration and XY sex chromosome complement, and negative responses to increasing estradiol concentration. While this appears to contradict the literature showing upregulation of the interferon pathways with estradiol and XX sex chromosome complement, our study represents the baseline, healthy-state, whereas the literature typically focuses on the female sex-bias after immunogenic stimulation, which represents the disease-state. In the future, investigating the roles of the sex hormones and sex chromosomes in health and through the transition into disease could provide further mechanistic insights into the sex-differential nature of autoimmunity and a plethora of other diseases.

## Acknowledgements

We acknowledge and thank members of the Page Lab, including Tatyana Pyntikova and Natalia Koutseva for sample processing, Linyong Mao for analysis guidance, Winston Bellott for comments on the manuscript, and Neha Bokil for assistance with visualization. We thank Boston Children’s Hospital employees Ashley Bass and Mina Wilcha for assistance with recruitment and Evan Schafer for sample processing. We thank the Genome Technology Core at the Whitehead Institute for library preparation and sequencing. We thank the University of Virginia Ligand Assay & Analysis Core for assistance with sex hormone assays. We thank the individuals who contributed samples for their participation.

## Author contributions

Conceptualization: RMH, YMC, and DCP. Methodology: RMH, TW, LVB, LM, and HS. Software: RMH, TW, LVB, LM, and HS. Validation: LVB and HS. Formal analysis: RMH, TW, LVB, LM, HS, and EJ. Investigation: RMH, TW, and LVB. Resources: RMH, KB, PH, EM, KS, PT, BL, and YMC. Data curation: RMH, LVB, and TW. Writing - original draft preparation: RMH and DCP. Writing - review and editing: RMH, TW, LVB, HS, JFH, and DCP. Visualization: RMH, TW, and LVB. Supervision: RMH and DCP. Project administration: RMH and DCP. Funding acquisition: RMH and DCP.

## Supplemental information titles and legends

Document S1. Figures S1-S2 and Tables S1 and S4.

Table S2. Responses of all significant autosomal genes to testosterone, estradiol, or sex chromosome complement, related to Figure 3.

Table S3. Mean-normalized pseudobulk counts for all autosomal genes with non-zero counts in at least 10% of the cells for the given PBMC cell type, related to Figure 3.

Table S5. Responses of all significant X-chromosomal genes to testosterone, estradiol, or sex chromosome complement, related to Figure 5.

Table S6. Mean-normalized pseudobulk counts for all X-chromosomal genes with non-zero counts in at least 10% of the cells for the given PBMC cell type, related to Figure 5.

## Methods

### Cohort recruitment

Between July 15^th^, 2019 and April 22^nd^, 2022, we enrolled 69 individuals for our research study. Our study was approved by the institutional review board at Boston Children’s Hospital (IRB-P00029637). We recruited 34 transgender individuals from the Gender Multispecialty Service (GeMS) clinic at Boston Children’s Hospital. In order to participate in this study, transgender individuals must have received exogenous estradiol or testosterone for at least a year as part of their medical care. We recruited 35 cisgender individuals from the general population. Cisgender individuals were required to be between the ages of 14-30 years. Exclusion criteria for all individuals were pregnancy or inability to provide consent. Cisgender individuals were also excluded if they had any medical diagnoses or used any prescription medications. We obtained written informed consent from each individual. If the individual was a minor, we obtained written informed assent from the individual and written informed consent from a guardian. Individuals completed a brief demographic questionnaire. We obtained body mass index (BMI) from the electronic medical record for transgender individuals if they had a clinical visit the same day the research study visit was completed. For the remaining individuals, we calculated BMI based on height and weight measurements obtained at the research study visit. We calculated age based on each individual’s report of birth date and we confirmed age from the electronic medical record for transgender individuals.

### Sample collection and processing

We obtained blood samples from each individual in an ethylenediaminetetraacetic acid (EDTA) tube (BD Vacutainer cat. #366643) and a serum-separating tube (SST) (BD Vacutainer cat. #367986). Samples were couriered to the Page laboratory at room temperature for same day processing. We isolated PBMCs using the 10x Genomics Sample Preparation Demonstrated Protocol: Fresh Frozen Human Peripheral Blood Mononuclear Cells for Single Cell RNA Sequencing (CG00039 Rev C). We placed up to 15 mL of blood from the EDTA tube in a new 50 mL conical tube, added PBS to a final volume of 40 ml, and gently inverted the tube. We added 10.5 mL of lymphocyte separation media (MP Biomedicals cat. #50494) to the tube to create a gradient. We spun the tubes at 1500 rpm (acceleration 5; brake 1) for 30 minutes at room temperature. We isolated PBMCs from the buffy coat and transferred them to a new 50 mL conical tube and we added PBS to a final volume of 40 mL. We centrifuged tubes at 1500 rpm for 10 minutes at room temperature. We removed the supernatant and resuspended the pellet in 5 mL PBS and we counted cells using the Countess II Automated Cell Counter via standard protocol from the manufacturer. We spun the tubes at 1500 rpm for 5 minutes at room temperature. We removed the PBS and resuspended the PBMCs in freezing media (FBS + 10% DMSO). We aliquoted the cells into cryovials and froze them in a styrofoam case at −80°C for at least 24 hours and then transferred them to liquid nitrogen.

For serum isolation, we gently inverted the SST and transferred the blood sample to a new tube. We centrifuged the tube at 2500 rpm for 15 minutes at 4°C. We isolated the serum and stored it in cryovials at −80°C.

### Serum hormone measurements and calculation of free hormone concentration

We shipped serum on dry ice to the University of Virginia Center for Research in Reproduction Ligand Assay & Analysis Core. All measurements were assayed using commercially available kits and standard protocols. The core assayed stradiol using the MP Biomedicals radioimmunoassay kit (catalog #07-238102); testosterone and sex hormone binding globulin (SHBG) using the Siemens Healthcare Diagnostics Immulite 2000 (testosterone catalog #L2KTW2/10381190; SHBG catalog #L2KSH2/10381198). Steroid assay validation procedures were based on recommendations from the Endocrine Society Council “Sex Steroid Assays Reporting Task Force.”^109^ We calculated free (“bioavailable”) testosterone and estradiol levels using total levels of testosterone or estradiol and SHBG via the mass action method of Vermeulen.^66^

### 10X 3’ RNA-sequencing

We thawed PBMCs according to the 10x Genomics Sample Preparation Demonstrated Protocol: Fresh Frozen Human Peripheral Blood Mononuclear Cells for Single Cell RNA Sequencing (CG00039 Rev C). We thawed PBMCs from up to four individuals at a time. To mitigate batch effects between the groups, we did not thaw PBMCs from individuals from the same group at the same time.

We removed a cryovial containing PBMCs from liquid nitrogen and transported it on dry ice to a water bath. We thawed the vial in a 37°C water bath for 2-3 minutes. We transferred the cell suspension to a tube and added 32 mL of media (RPMI + 10% FBS) gradually. We centrifuged the tube at 300 rcf for 5 minutes at 22°C (acceleration 5; break 1). We discarded the supernatant and resuspended the cells in the remaining 1 mL of media. We added an additional 9 mL of media. We counted the cells using the Countess II Automated Cell Counter via the manufacturer’s standard protocol. We transferred 2 million cells to a new tube and centrifuged it at 300 rcf for 5 minutes at 22°C (acceleration 5; break 1). We removed the supernatant and gently resuspended the cells in 1 mL PBS + 0.04% BSA, and transferred the solution to a 2 mL DNA LoBind tube (Eppendorf cat. #022431048). We rinsed the tube with 0.5 mL PBS + 0.04% BSA and transferred the volume to the same Eppendorf tube. We centrifuged the Eppendorf tube at 300 rcf for 5 minutes at 22°C. We removed the supernatant and gently resuspended the pellet in 2 mL PBS + 0.04% BSA until a single cell suspension was achieved. We counted the cells using the Countess II Automated Cell Counter via the manufacturer’s standard protocol.

We brought 70,000-120,000 PBMCs/sample on ice to the Genome Technology Core at the Whitehead Institute. The core processed the cells using the 10X Genomics Chromium Controller with the Next GEM Single Cell 3’ v3.1 Reagent Kit, according to manufacturer’s directions. The core achieved a target of approximately 10,000 cells per library by loading 1.65 times the target number of cells in suspension along with barcoded beads and partitioning oil into the Chromium Controller, in order to create gel beads in emulsion. The Chromium Controller combined individual cells, first strand master mix, and gel beads containing barcoded oligonucleotides into single-cell droplets for first strand cDNA synthesis, so that each cell was marked with its own unique barcode during reverse transcription. The 3’ beads contained a poly(dT) oligo that enabled the production of barcoded, full-length cDNA from poly-adenylated mRNA. After first strand synthesis was complete, the core dissolved the emulsion and pooled the cDNA for bulk processing as a single sample. The core fragmented, end-repaired, A-tailed and ligated the sample with universal adapters. The core added a second sample barcode during the PCR step, allowing for unique library identification. Thus, the core created a single library containing data for each individual cell. The core sequenced samples using a NovaSeq 6000 S4 flow cell with a read length of 150×150 base pairs. The core completed quality-control analysis using the ThermoFisher Scientific Qubit Fluorometric Quantification 3 and the Agilent 2100 Bioanalyzer.

### scRNA-seq data processing and analysis

We performed all analyses using the human reference genome (GRCh38), and a custom version of the comprehensive GENCODE, release 24 transcriptome annotation.^110^ This annotation represents the union of the ‘‘GENCODE Basic’’ annotation and transcripts recognized by the Consensus Coding Sequence project. Importantly, the GENCODE annotation lists the pseudoautosomal (PAR) gene annotations twice, once on the X chromosome and once on the Y chromosome, which complicates analysis. We removed these annotations from the Y chromosome so the PAR genes are only listed once in our annotation, on the X chromosome.

We pseudoaligned reads to the human transcriptome and estimated transcript counts using the kallisto and bustools programs (v.0.46.1).^111,112^ We analyzed the estimated counts using the Seurat package (v. 4.3.0) in R (v. 4.2.1).^113,114^ We retained cells for further analysis if they had at least 500 features per cell, less than 15% mitochondrial gene expression,^115^ and a binary classification based doublet score less than 0.8 using the scds program.^116^ We identified subtypes of PBMCs using marker genes (Table S1). For the rare HSPC and plasmablast cell subtypes, we assigned cell types via alignment to a single-cell PBMC reference atlas.^73^ For each cell type, we assessed differential gene expression using DESeq2 (v. 1.36.0)^117^ with a pseudobulk approach.^118^ For a given cell type, we tested genes for differential expression only if they exhibited non-zero counts in at least 10% of the cells. Within the negative binomial framework of DESeq2, we modeled gene expression by an additive combination of the following covariates: calculated free testosterone, calculated free estradiol, sex chromosome complement, BMI, age, and batch. We included sex chromosome complement and batch as categorical covariates and the remaining covariates as continuous. We mean-centered gene expressed. We carried out gene-set enrichment analysis (GSEA) on the Hallmark collection of gene-sets from the MSigDB^87^ using the fgsea package (v. 1.22.0)^119^ for R with genes pre-ranked according to the Wald statistic from DESeq2.^86^

### Statistical analysis

We used a variety of statistical tests to calculate *p*-values, which are noted in the manuscript text or figure legends as appropriate. We used R software version 3.6.3 to calculate statistics and generate plots.^120^ We considered results statistically significant when *p*<0.05, or adjusted-*p* or FDR<0.05 when multiple hypothesis correction was applied. Data are shown as median and interquartile range, unless stated otherwise. We created the Venn diagrams using BioVenn.^121^

## References

1. Gray JP, Wolfe LD. Height and sexual dimorphism of stature among human societies. Am J Phys Anthropol. 1980;53(3):441–456. doi:10.1002/ajpa.1330530314

2. Brix N, Ernst A, Lauridsen LLB, et al. Timing of puberty in boys and girls: A population-based study. Paediatr Perinat Epidemiol. 2019;33(1):70–78. doi:10.1111/ppe.12507

3. Regitz-Zagrosek V. Sex and gender differences in health. EMBO Rep. 2012;13(7):596–603. doi:10.1038/embor.2012.87

4. Morrow EH. The evolution of sex differences in disease. Biol Sex Differ. 2015;6(1):1–7. doi:10.1186/s13293-015-0023-0

5. Ober C, Loisel DA, Gilad Y. Sex-specific genetic architecture of human disease. Nat Rev Genet. 2008;9(12):911–922. doi:10.1038/nrg2415

6. Heron M. Deaths: Leading causes for 2017. National Vital Statistics Reports. 2019;68(6):1–76.

7. Mauvais-Jarvis F, Bairey Merz N, Barnes PJ, et al. Sex and gender: modifiers of health, disease, and medicine. The Lancet. 2020;396(10250):565–582. doi:10.1016/S0140-6736(20)31561-0

8. Battey R. Normal ovariotomectomy. Atlanta Medical and Surgical Journal. 1873;10:321–339.

9. Berthold A. Transplantation der hoden. Arch Anat Physiol Wissensch. Published online 1849:42.

10. Knauer E. Transplantation of the ovaries: an experimental study. Archiv F Gyn. Published online 1898.

11. Knauer E. On ovarian transplantation: labour and normal end of pregnancy after transplantation of ovaries in rabbits. Contralb F Gyn. 1898;8.

12. Knauer E. Ovarian transplantation in rabbits: a preliminary report. Contralb F Gyn. Published online 1898.

13. Hunter J. Treaties on the Natural History of the Human Teeth. In: Palmer J, ed. The Works of John Hunter F.R.S. Vol 2. Longman, Rees, Orme, Brown, Green, and Longman; 1835:56.

14. Fosbery W. Severe climacteric flushings successfully treated with ovarian extract. BMJ. 1897;1:1039.

15. Brown-Séquard. Note on the effects produced on man by subcutaneous injections of a liquid obtained from the testicles of animals. The Lancet. 1889;134(3438). doi:10.1016/S0140-6736(00)64118-1

16. Steinach E. Verjüngung Durch Experimentelle Neubelebung Der Alternden. Julius Springer; 1920.

17. Sturgis SH, Albright F. The mechanism of estrin therapy in the relief of dysmenorrhea. Endocrinology. 1940;26(1). doi:10.1210/endo-26-1-68

18. Albright F. Studies on ovarian dysfunction. III. The menopause. Endocrinology. 1936;20(1). doi:10.1210/endo-20-1-24

19. Albright F, Halsted JA, Cloney E. Studies on ovarian dysfunction. New England Journal of Medicine. 1935;212(5). doi:10.1056/nejm193501312120503

20. Bender CM, Merriman JD, Gentry AL, et al. Patterns of change in cognitive function with anastrozole therapy. Cancer. 2015;121(15). doi:10.1002/cncr.29393

21. Kim JK, Levin ER. Estrogen signaling in the cardiovascular system. Nucl Recept Signal. 2006;4(1). doi:10.1621/nrs.04013

22. Chambliss KL, Wu Q, Oltmann S, et al. Non-nuclear estrogen receptor α signaling promotes cardiovascular protection but not uterine or breast cancer growth in mice. Journal of Clinical Investigation. 2010;120(7). doi:10.1172/JCI38291

23. Mendelsohn ME, Karas RH. HRT and the Young at Heart. New England Journal of Medicine. 2007;356(25). doi:10.1056/nejme078072

24. Finkelstein JS, Lee H, Burnett-Bowie SAM, et al. Gonadal Steroids and Body Composition, Strength, and Sexual Function in Men. New England Journal of Medicine. 2013;369(11). doi:10.1056/nejmoa1206168

25. Simpson ER, Brown KA. Minireview: Obesity and breast cancer: A Tale of inflammation and dysregulated metabolism. Molecular Endocrinology. 2013;27(5). doi:10.1210/me.2013-1011

26. Ohno S. Sex Chromosomes and Sex-Linked Genes. In: Monographs on Endocrinology. Vol 1.; 1966.

27. Lahn BT, Page DC. Four evolutionary strata on the human X chromosome. Science (1979). 1999;286(5441). doi:10.1126/science.286.5441.964

28. Lyon MF. Gene action in the X-chromosome of the mouse (mus musculus L.). Nature. 1961;190(4773). doi:10.1038/190372a0

29. Brown CJ, Ballabio A, Rupert JL, et al. A gene from the region of the human X inactivation centre is expressed exclusively from the inactive X chromosome. Nature. 1991;349(6304). doi:10.1038/349038a0

30. Stern C. The problem of complete Y-linkage in man. Am J Hum Genet. 1957;9(3).

31. Skaletsky H, Kuroda-Kawaguchl T, Minx PJ, et al. The male-specific region of the human Y chromosome is a mosaic of discrete sequence classes. Nature. 2003;423(6942). doi:10.1038/nature01722

32. Carrel L, Willard HF. X-inactivation profile reveals extensive variability in X-linked gene expression in females. Nature. 2005;434(7031). doi:10.1038/nature03479

33. Tukiainen T, Villani AC, Yen A, et al. Landscape of X chromosome inactivation across human tissues. Nature. 2017;550(7675):244–248. doi:10.1038/nature24265

34. Balaton BP, Cotton AM, Brown CJ. Derivation of consensus inactivation status for X-linked genes from genome-wide studies. Biol Sex Differ. 2015;6(1). doi:10.1186/s13293-015-0053-7

35. Cotton AM, Ge B, Light N, Adoue V, Pastinen T, Brown CJ. Analysis of expressed SNPs identifies variable extents of expression from the human inactive X chromosome. Genome Biol. 2013;14(11). doi:10.1186/gb-2013-14-11-r122

36. Garieri M, Stamoulis G, Blanc X, et al. Extensive cellular heterogeneity of X inactivation revealed by single-cell allele-specific expression in human fibroblasts. Proc Natl Acad Sci U S A. 2018;115(51). doi:10.1073/pnas.1806811115

37. Wainer Katsir K, Linial M. Human genes escaping X-inactivation revealed by single cell expression data. BMC Genomics. 2019;20(1). doi:10.1186/s12864-019-5507-6

38. Sauteraud R, Stahl JM, James J, et al. Inferring genes that escape X-Chromosome inactivation reveals important contribution of variable escape genes to sex-biased diseases. Genome Res. 2021;31(9). doi:10.1101/gr.275677.121

39. San Roman AK, Godfrey AK, Skaletsky H, et al. The human inactive X chromosome modulates expression of the active X chromosome. Cell Genomics. 2023;3(2). doi:10.1016/j.xgen.2023.100259

40. Bellott DW, Hughes JF, Skaletsky H, et al. Mammalian y chromosomes retain widely expressed dosage-sensitive regulators. Nature. 2014;508(7497):494-499. doi:10.1038/nature13206

41. San Roman AK, Skaletsky H, Godfrey AK, et al. The human Y and inactive X chromosomes similarly modulate autosomal gene expression. Cell Genomics. Published online 2023. doi:10.1016/j.xgen.2023.100462

42. Dragin N, Bismuth J, Cizeron-Clairac G, et al. Estrogen-mediated downregulation of AIRE influences sexual dimorphism in autoimmune diseases. Journal of Clinical Investigation. 2016;126(4):1525–1537. doi:10.1172/JCI81894

43. Naqvi S, Godfrey AK, Hughes JF, Goodheart ML, Mitchell RN, Page DC. Conservation, acquisition, and functional impact of sex-biased gene expression in mammals. Science (1979). 2019;365(6450):eaaw7317. doi:10.1126/science.aaw7317

44. Aguet F, Barbeira AN, Bonazzola R, et al. The impact of sex on gene expression across human tissues. Science (1979). 2020;369(6509). doi:10.1126/SCIENCE.ABA3066

45. Lopes-Ramos CM, Chen CY, Kuijjer ML, et al. Sex Differences in Gene Expression and Regulatory Networks across 29 Human Tissues. Cell Rep. 2020;31(12):107795. doi:10.1016/j.celrep.2020.107795

46. Klein SL, Flanagan KL. Sex differences in immune responses. Nat Rev Immunol. 2016;16:626–638.

47. Potluri T, Fink AL, Sylvia KE, et al. Age-associated changes in the impact of sex steroids on influenza vaccine responses in males and females. NPJ Vaccines. 2019;4(1). doi:10.1038/s41541-019-0124-6

48. Shapiro JR, Morgan R, Leng SX, Klein SL. Roadmap for Sex-Responsive Influenza and COVID-19 Vaccine Research in Older Adults. Frontiers in Aging. 2022;3. doi:10.3389/fragi.2022.836642

49. Fink AL, Klein SL. The evolution of greater humoral immunity in females than males: implications for vaccine efficacy. Curr Opin Physiol. 2018;6. doi:10.1016/j.cophys.2018.03.010

50. Klein SL, Jedlicka A, Pekosz A. The Xs and Y of immune responses to viral vaccines. Lancet Infect Dis. 2010;10(5). doi:10.1016/S1473-3099(10)70049-9

51. Flanagan KL, Fink AL, Plebanski M, Klein SL. Sex and gender differences in the outcomes of vaccination over the life course. Annu Rev Cell Dev Biol. 2017;33. doi:10.1146/annurev-cellbio-100616-060718

52. Klein SL. Sex influences immune responses to viruses, and efficacy of prophylaxis and treatments for viral diseases. BioEssays. 2012;34(12). doi:10.1002/bies.201200099

53. Ursin RL, Shapiro JR, Klein SL. Sex-biased Immune Responses Following SARS-CoV-2 Infection. Trends Microbiol. 2020;28(12). doi:10.1016/j.tim.2020.10.002

54. Scully EP, Haverfield J, Ursin RL, Tannenbaum C, Klein SL. Considering how biological sex impacts immune responses and COVID-19 outcomes. Nat Rev Immunol. 2020;20(7). doi:10.1038/s41577-020-0348-8

55. Klein SL, Passaretti C, Anker M, Olukoya P, Pekosz A. The impact of sex, gender and pregnancy on 2009 H1N1 disease. Biol Sex Differ. 2010;1(1). doi:10.1186/2042-6410-1-5

56. Klein SL. Hormonal and immunological mechanisms mediating sex differences in parasite infection. Parasite Immunol. 2004;26(6-7). doi:10.1111/j.0141-9838.2004.00710.x

57. Ursin RL, Klein SL. Sex Differences in Respiratory Viral Pathogenesis and Treatments. Annu Rev Virol. 2021;8. doi:10.1146/annurev-virology-091919-092720

58. Whitacre C. Sex differences in autoimmune disease. Nat Immunol. 2001;2:777–780.

59. Lin X, Yu H, Zhao C, et al. The Peripheral Blood Mononuclear Cell Count Is Associated With Bone Health in Elderly Men: A Cross-Sectional Population-Based Study. Medicine. 2016;95(15):e3357. doi:10.1097/MD.0000000000003357

60. Bahr TM, Hughes GJ, Armstrong M, et al. Peripheral blood mononuclear cell gene expression in chronic obstructive pulmonary disease. Am J Respir Cell Mol Biol. 2013;49(2):316–323. doi:10.1165/rcmb.2012-0230OC

61. Lassale C, Curtis A, Abete I, et al. Elements of the complete blood count associated with cardiovascular disease incidence: Findings from the EPIC-NL cohort study. Sci Rep. 2018;8(3290):1-11. doi:10.1038/s41598-018-21661-x

62. Pan C, Kumar C, Bohl S, Klingmueller U, Mann M. Comparative Proteomic Phenotyping of Cell Lines and Primary Cells to Assess Preservation of Cell Type-specific Functions. Molecular & Cellular Proteomics. 2009;8(3):443–450. doi:10.1074/mcp.M800258-MCP200

63. Tischfield JA, Mbarek H, Milaneschi Y, et al. Sex differences in the human peripheral blood transcriptome. BMC Genomics. 2014;15(1):33. doi:10.1186/1471-2164-15-33

64. Ghanim H, Dhindsa S, Abuaysheh S, et al. Diminished androgen and estrogen receptors and aromatase levels in hypogonadal diabetic men: Reversal with testosterone. Eur J Endocrinol. 2018;178(3):277–283. doi:10.1530/EJE-17-0673

65. Scariano JK, Emery-Cohen AJ, Pickett GG, Morgan M, Simons PC, Alba F. Estrogen receptors alpha (ESR1) and beta (ESR2) are expressed in circulating human lymphocytes. Journal of Receptors and Signal Transduction. 2008;28(3):285–293. doi:10.1080/10799890802084614

66. Vermeulen A, Verdonck L, Kaufman JM. A critical evaluation of simple methods for the estimation of free testosterone in serum. Journal of Clinical Endocrinology and Metabolism. 1999;84(10). doi:10.1210/jcem.84.10.6079

67. Huang Z, Chen B, Liu X, et al. Effects of sex and aging on the immune cell landscape as assessed by single-cell transcriptomic analysis. Proc Natl Acad Sci U S A. 2021;118(33). doi:10.1073/pnas.2023216118

68. Peters MJ, Joehanes R, Pilling LC, et al. The transcriptional landscape of age in human peripheral blood. Nat Commun. 2015;6. doi:10.1038/ncomms9570

69. Zheng Y, Liu X, Le W, et al. A human circulating immune cell landscape in aging and COVID-19. Protein Cell. 2020;11(10). doi:10.1007/s13238-020-00762-2

70. Liu N, Chen X, Ran J, et al. Investigating the change in gene expression profile of blood mononuclear cells post-laparoscopic sleeve gastrectomy in Chinese obese patients. Front Endocrinol (Lausanne). 2023;14. doi:10.3389/fendo.2023.1049484

71. Costa A, van der Stelt I, Reynés B, et al. Whole-Genome Transcriptomics of PBMC to Identify Biomarkers of Early Metabolic Risk in Apparently Healthy People with Overweight-Obesity and in Normal-Weight Subjects. Mol Nutr Food Res. 2023;67(4). doi:10.1002/mnfr.202200503

72. Jang K, Tong T, Lee J, Park T, Lee H. Altered Gene Expression Profiles in Peripheral Blood Mononuclear Cells in Obese Subjects. Obes Facts. 2020;13(3). doi:10.1159/000507817

73. Hao Y, Hao S, Andersen-Nissen E, et al. Integrated analysis of multimodal single-cell data. Cell. 2021;184(13):3573–3587.e29. doi:10.1016/j.cell.2021.04.048

74. Márquez EJ, Chung C han, Marches R, et al. Sexual-dimorphism in human immune system aging. Nat Commun. 2020;11(1). doi:10.1038/s41467-020-14396-9

75. Gagliano-Jucá T, Pencina KM, Guo W, et al. Differential effects of testosterone on circulating neutrophils, monocytes, and platelets in men: Findings from two trials. Andrology. 2020;8(5). doi:10.1111/andr.12834

76. Olsen NJ, Kovacs WJ. Evidence that androgens modulate human thymic T cell output. Journal of Investigative Medicine. 2011;59(1). doi:10.2310/JIM.0b013e318200dc98

77. Sutherland JS, Goldberg GL, Hammett M V., et al. Activation of Thymic Regeneration in Mice and Humans following Androgen Blockade. The Journal of Immunology. 2005;175(4). doi:10.4049/jimmunol.175.4.2741

78. Blanton L V, San Roman AK, Wood G, et al. Stable and robust Xi and Y transcriptomes drive cell-type-specific autosomal and Xa responses in vivo and in vitro in four human cell types. Cell Genomics. 2024;4:1–16.

79. Kory N, Wyant GA, Prakash G, et al. SFXN1 is a mitochondrial serine transporter required for one-carbon metabolism. Science (1979). 2018;362(6416). doi:10.1126/science.aat9528

80. Ducker GS, Rabinowitz JD. One-Carbon Metabolism in Health and Disease. Cell Metab. 2017;25(1). doi:10.1016/j.cmet.2016.08.009

81. Locasale JW. Serine, glycine and one-carbon units: Cancer metabolism in full circle. Nat Rev Cancer. 2013;13(8). doi:10.1038/nrc3557

82. Yang M, Vousden KH. Serine and one-carbon metabolism in cancer. Nat Rev Cancer. 2016;16(10). doi:10.1038/nrc.2016.81

83. Kim R, Frederik Nijhout H, Reed MC. One-carbon metabolism during the menstrual cycle and pregnancy. PLoS Comput Biol. 2021;17(12). doi:10.1371/journal.pcbi.1009708

84. Sadre-Marandi F, Dahdoul T, Reed MC, Nijhout HF. Sex differences in hepatic one-carbon metabolism. BMC Syst Biol. 2018;12(1). doi:10.1186/s12918-018-0621-7

85. Howe CG, Liu X, Hall MN, et al. Sex-specific associations between one-carbon metabolism indices and posttranslational histone modifications in arsenic-exposed Bangladeshi adults. Cancer Epidemiology Biomarkers and Prevention. 2017;26(2). doi:10.1158/1055-9965.EPI-16-0202

86. Subramanian A, Tamayo P, Mootha VK, et al. Gene set enrichment analysis: A knowledge-based approach for interpreting genome-wide expression profiles. Proc Natl Acad Sci U S A. 2005;102(43):15545–15550. doi:10.1073/pnas.0506580102

87. Liberzon A, Birger C, Thorvaldsdottir H, Ghandi M, Mesirov JP, Tamayo P. The Molecular Signatures Database (MSigDB) hallmark gene set collection. Cell Syst. 2015;1(6):417–425. doi:10.1016/j.cels.2015.12.004

88. Berghöfer B, Frommer T, Haley G, Fink L, Bein G, Hackstein H. TLR7 Ligands Induce Higher IFN-α Production in Females. The Journal of Immunology. 2006;177(4). doi:10.4049/jimmunol.177.4.2088

89. Zhang MA, Rego D, Moshkova M, et al. Peroxisome proliferator-activated receptor (PPAR)α and -γ regulate IFNγ and IL-17A production by human T cells in a sex-specific way. Proc Natl Acad Sci U S A. 2012;109(24):9505–9510. doi:10.1073/pnas.1118458109

90. Crow MK. NEWS & VIEWS Interferon α or β: which is the culprit in autoimmune disease ? Nature Publishing Group. 2016;12(8):439–440. doi:10.1038/nrrheum.2016.117

91. Pelfrey CM, Cotleur AC, Lee JC, Rudick RA. Sex differences in cytokine responses to myelin peptides in multiple sclerosis. J Neuroimmunol. 2002;130(1-2). doi:10.1016/S0165-5728(02)00224-2

92. Ivashkiv LB, Donlin LT. Regulation of type i interferon responses. Nat Rev Immunol. 2014;14(1). doi:10.1038/nri3581

93. Rubtsova K, Marrack P, Rubtsov A V. TLR7, IFNc, and T-bet: Their roles in the development of ABCs in female-biased autoimmunity. Cell Immunol. 2015;294(2). doi:10.1016/j.cellimm.2014.12.002

94. Krongrad A, Wilson CM, Wilson JD, Allman DR, McPhaul MJ. Androgen increases androgen receptor protein while decreasing receptor mRNA in LNCaP cells. Mol Cell Endocrinol. 1991;76(1-3). doi:10.1016/0303-7207(91)90262-Q

95. Wolf DA, Herzinger T, Hermeking H, Blaschke D, Hörz W. Transcriptional and posttranscriptional regulation of human androgen receptor expression by androgen. Molecular Endocrinology. 1993;7(7). doi:10.1210/mend.7.7.8413317

96. Wiren KM, Zhang X, Chang C, Keenan E, Orwoll ES. Transcriptional up-regulation of the human androgen receptor by androgen in bone cells. Endocrinology. 1997;138(6). doi:10.1210/endo.138.6.5163

97. Smeds L, Kojola I, Ellegren H. The evolutionary history of grey wolf Y chromosomes. Mol Ecol. 2019;28(9). doi:10.1111/mec.15054

98. Auer MK, Cecil A, Roepke Y, et al. 12-months metabolic changes among gender dysphoric individuals under cross-sex hormone treatment: A targeted metabolomics study. Sci Rep. 2016;6(November):1–10. doi:10.1038/srep37005

99. Shepherd R, Bretherton I, Pang K, et al. Gender-affirming hormone therapy induces specific DNA methylation changes in blood. Clin Epigenetics. 2022;14(1):1–19. doi:10.1186/s13148-022-01236-4

100. Robinson GA, Peng J, Peckham H, et al. Investigating sex differences in T regulatory cells from cisgender and transgender healthy individuals and patients with autoimmune inflammatory disease: a cross-sectional study. Lancet Rheumatol. 2022;4(10):e710–e724. doi:10.1016/S2665-9913(22)00198-9

101. Grünhagel B, Borggrewe M, Hagen SH, et al. Reduction of IFN-I responses by plasmacytoid dendritic cells in a longitudinal trans men cohort. iScience. 2023;26(11). doi:10.1016/j.isci.2023.108209

102. Maru J, Millington K, Carswell J. Greater Than Expected Prevalence of Type 1 Diabetes Mellitus Found in an Urban Gender Program. Transgend Health. 2021;6(1):57–60. doi:10.1089/trgh.2020.0027

103. Logel SN, Bekx MT, Rehm JL. Potential association between type 1 diabetes mellitus and gender dysphoria. Pediatr Diabetes. 2020;21(2). doi:10.1111/pedi.12947

104. Logel S, Maru J, Whitehead J, et al. Higher Rates of Certain Autoimmune Diseases in Transgender and Gender Diverse Youth. Transgend Health. Published online 2023.

105. Defreyne J, De Bacquer D, Shadid S, Lapauw B, T’Sjoen G. Is Type 1 Diabetes Mellitus More Prevalent Than Expected in Transgender Persons? A Local Observation. Sex Med. 2017;5(3):e215–e218. doi:10.1016/j.esxm.2017.06.004

106. Campochiaro C, Host L V., Ong VH, Denton CP. Development of systemic sclerosis in transgender females: A case series and review of the literature. Clin Exp Rheumatol. 2018;36.

107. Chan KL, Mok CC. Development of systemic lupus erythematosus in a male-to-female transsexual: The role of sex hormones revisited. Lupus. 2013;22(13). doi:10.1177/0961203313500550

108. Hill BG, Hodge B, Misischia R. Lupus nephritis in a transgender woman on cross-sex hormone therapy: a case for the role of oestrogen in systemic lupus erythematosus. Lupus. 2020;29(13). doi:10.1177/0961203320946372

109. The New Instructions to Authors for the Reporting of Steroid Hormone Measurements. Horm Cancer. 2014;5(6). doi:10.1007/s12672-014-0202-1

110. Godfrey AK, Naqvi S, Chmátal L, et al. Quantitative analysis of Y-Chromosome gene expression across 36 human tissues. Genome Res. Published online 2020:1–14. doi:10.1101/gr.261248.120

111. Melsted P, Sina Booeshaghi A, Gao F, et al. Modular and efficient pre-processing of single-cell RNA-seq. bioRxiv. Published online 2019.

112. Bray NL, Pimentel H, Melsted P, Pachter L. Near-optimal probabilistic RNA-seq quantification. Nat Biotechnol. 2016;34(5). doi:10.1038/nbt.3519

113. Team RC. R: A language and environment for statistical computing v. 3.6. 1 (R Foundation for Statistical Computing, Vienna, Austria, 2019). Scientific Reports.

114. Satija R, Farrell JA, Gennert D, Schier AF, Regev A. Spatial reconstruction of single-cell gene expression data. Nat Biotechnol. 2015;33(5). doi:10.1038/nbt.3192

115. Osorio D, Cai JJ. Systematic determination of the mitochondrial proportion in human and mice tissues for single-cell RNA sequencing data quality control. bioRxiv. Published online 2020. doi:10.1101/2020.02.20.958793

116. Bais AS, Kostka D. Scds: Computational annotation of doublets in single-cell RNA sequencing data. Bioinformatics. 2020;36(4). doi:10.1093/bioinformatics/btz698

117. Love MI, Huber W, Anders S. Moderated estimation of fold change and dispersion for RNA-seq data with DESeq2. Genome Biol. 2014;15(12). doi:10.1186/s13059-014-0550-8

118. Squair JW, Gautier M, Kathe C, et al. Confronting false discoveries in single-cell differential expression. Nat Commun. 2021;12(1). doi:10.1038/s41467-021-25960-2

119. Korotkevich G, Sukhov V, Budin N, Atryomov MN, Sergushichev A. Fast gene set enrichment analysis. bioRxiv. bioRxiv. Published online 2021.

120. R Core Team (2021). R: A Language and Environment for Statistical Computing. R Foundation for Statistical Computing. Published online 2021.

121. Hulsen T, de Vlieg J, Alkema W. BioVenn - A web application for the comparison and visualization of biological lists using area-proportional Venn diagrams. BMC Genomics. 2008;9. doi:10.1186/1471-2164-9-488

