## Supplementary material for "Independent effects of testosterone, estradiol, and sex chromosomes on gene expression in immune cells of trans- and cisgender individuals": Document S1.pdf

Figure S1

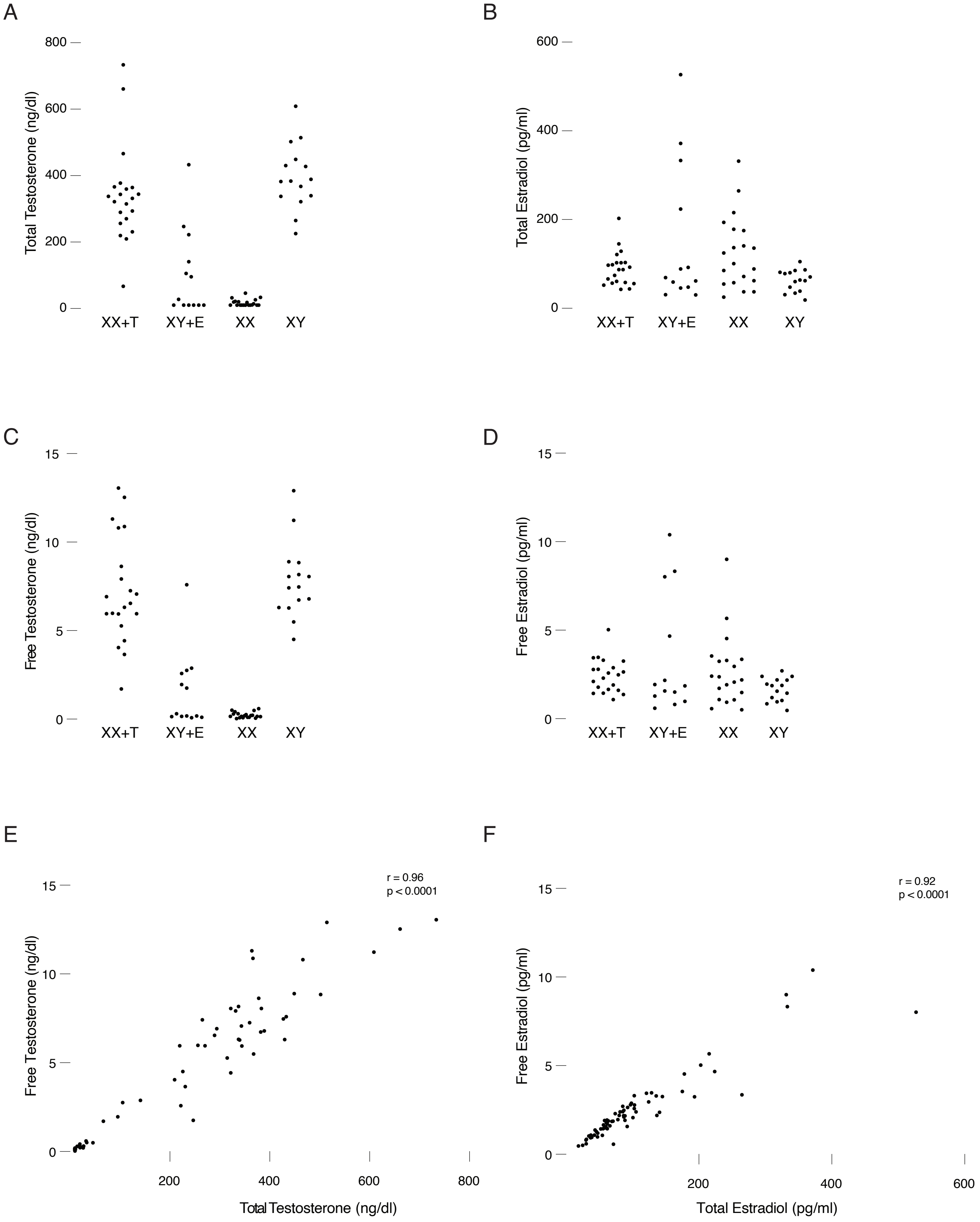

A
B
C
D
E
F
Total Testosterone (ng/dl)
Total Estradiol (pg/ml)
Free Testosterone (ng/dl)
Free Estradiol (pg/ml)
XX+T
XY+E
XX
XY
r = 0.96
p < 0.0001
r = 0.92
p < 0.0001
Total Testosterone (ng/dl)
Total Estradiol (pg/ml)

**Figure S1. Total and calculated free testosterone and estradiol concentrations separated by group.** Total testosterone **(A)**, total estradiol **(B)**, free testosterone **(C)**, and free estradiol **(D)** concentrations in all 69 individuals separated by group. Each point is one individual. The free fractions of testosterone and estradiol were calculated using the mass action method of Vermeulen (total testosterone or estradiol concentration / sex hormone binding globulin concentration). Correlation of total and free testosterone **(E)** and estradiol **(F)** concentrations. Pearson correlation coefficients and *p*-values shown.

Figure S2

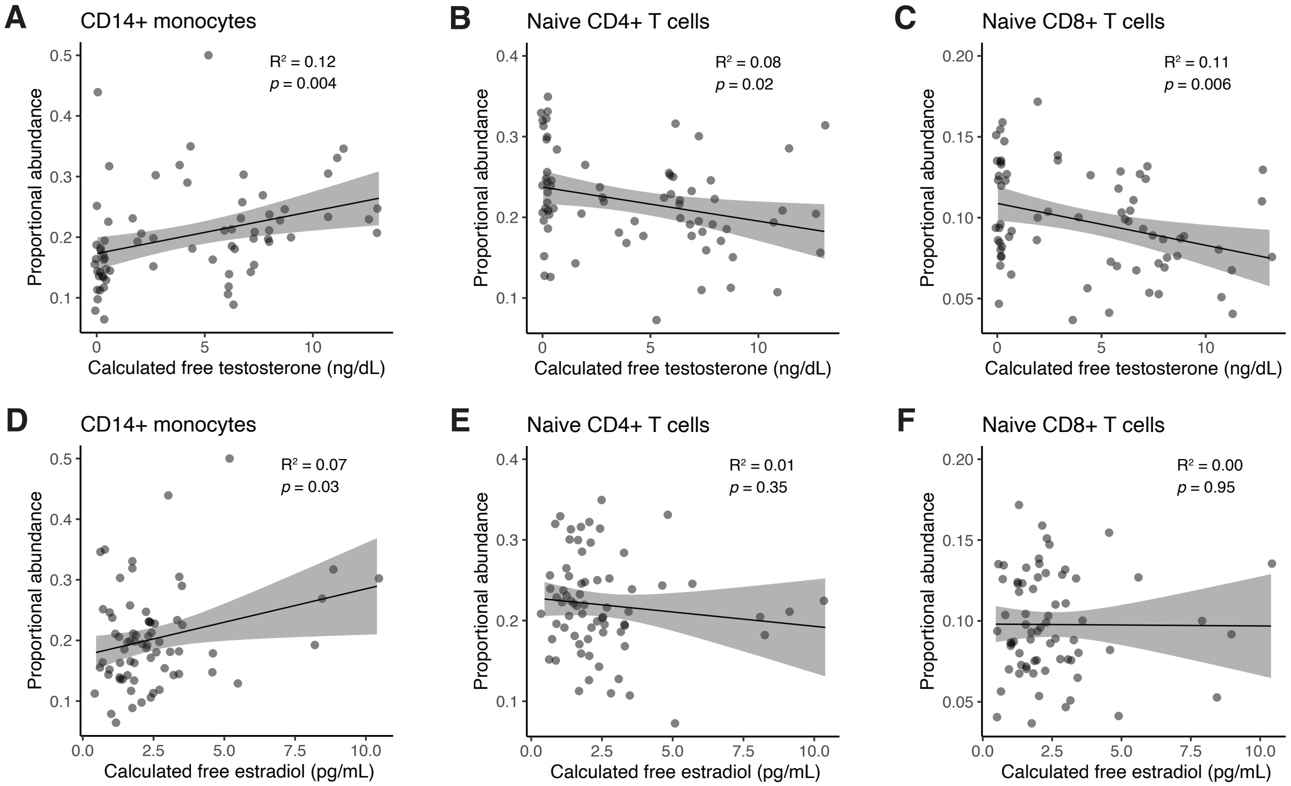


**Figure S2. Testosterone and estradiol effects on cell abundances.** Scatterplots of the relationship between calculated free testosterone or estradiol and proportional abundances of CD14+ monocytes **(A,D)**, naïve CD4+ T cells **(B,E)**, and naïve CD8+ T cells **(C,F)**. $R^2$ and *p*-values as shown.

| Cell type | Marker genes |
| --- | --- |
| CD4+ naïve T cells | CD3D+, CD4+, CCR7+ |
| CD4+ T cell memory ($T_{CM}$) | CD3D+, CD4+, CCR7+, SELL lo, ILR7A+ |
| CD8+ naïve T cells | CD3D+, CD8+, CCR7+, CD27+ |
| CD8+ T effector memory ($T_{EM}$) | CD3D+, CD8+, CCR7 lo, CD27 lo, IL7RA hi |
| CD8+ T effector memory cells re-expressing CD45RA ($T_{EMRA}$) | CD3D+, CD8+, CCR7-, CD27-, IL7RA lo, GZMB |
| Double negative T cells (DNT) | CD3D+, CD4-, CD8-, CD27+ |
| Regulatory T cells (Treg) | CD3D+, CD4+, CCR7-, FOXP3+ |
| Mucosal-associated invariant T cells (MAIT) | SLC4A10+ |
| CD56 bright natural killer (NK) | CD3D-, GNLY+, CD56 bright, SELL+ |
| CD56 dim natural killer (NK) | CD3D-, GNLY+, CD56 dim, SELL- |
| CD14+ monocytes | CD14+ |
| CD16+ monocytes | FCGR3A+ |
| Classical dendritic cells (cDC) | FCER1A+ |
| Plasmacytoid dendritic cells (pDC) | UGCG+ |
| B cells | MS4A1+ |

**Table S1. Marker genes for 15 peripheral blood mononuclear cell types, related to Figure 2.** Marker genes were selected for each cell type based on review of the literature.

| | CD14 | CD16 | cDC | pDC | HSPC | Plasmablast | B | CD4 naive | CD4 TCM | DNT | CD8 naive | CD8 TCM | CD8 TEM | CD8 TEMRA | Treg | MAIT | CD56 dim NK | CD56 bright NK |
|---|---|---|---|---|---|---|---|---|---|---|---|---|---|---|---|---|---|---|
| CD14 | 8052 | 7832 | 7924 | 7343 | 7590 | 7464 | 5469 | 4803 | 5596 | 6944 | 5297 | 7431 | 5341 | 5274 | 5511 | 5352 | 5531 | 5924 |
| CD16 | 7832 | 8475 | 8253 | 7606 | 7954 | 7868 | 5516 | 4820 | 5639 | 7182 | 5337 | 7804 | 5367 | 5300 | 5549 | 5378 | 5584 | 5983 |
| cDC | 7924 | 8253 | 9207 | 8062 | 8575 | 8443 | 5612 | 4897 | 5757 | 7478 | 5438 | 8297 | 5457 | 5377 | 5659 | 5477 | 5659 | 6093 |
| pDC | 7343 | 7606 | 8062 | 8423 | 8016 | 7922 | 5544 | 4834 | 5632 | 7195 | 5363 | 7789 | 5368 | 5269 | 5543 | 5371 | 5540 | 5967 |
| HSPC | 7590 | 7954 | 8575 | 8016 | 9512 | 8564 | 5589 | 4908 | 5753 | 7503 | 5456 | 8408 | 5465 | 5368 | 5658 | 5474 | 5664 | 6121 |
| Plasmablast | 7464 | 7868 | 8443 | 7922 | 8564 | 9364 | 5614 | 4878 | 5729 | 7486 | 5419 | 8565 | 5435 | 5334 | 5639 | 5447 | 5592 | 6028 |
| B | 5469 | 5516 | 5612 | 5544 | 5589 | 5614 | 5717 | 4696 | 5155 | 5485 | 5039 | 5578 | 5040 | 4902 | 5141 | 4964 | 5035 | 5280 |
| CD4 naive | 4803 | 4820 | 4897 | 4834 | 4908 | 4878 | 4696 | 5000 | 4960 | 4957 | 4959 | 4975 | 4882 | 4726 | 4939 | 4812 | 4720 | 4833 |
| CD4 TCM | 5596 | 5639 | 5757 | 5632 | 5753 | 5729 | 5155 | 4960 | 5893 | 5792 | 5384 | 5863 | 5460 | 5257 | 5609 | 5404 | 5312 | 5535 |
| DNT | 6944 | 7182 | 7478 | 7195 | 7503 | 7486 | 5485 | 4957 | 5792 | 7776 | 5499 | 7632 | 5517 | 5384 | 5714 | 5495 | 5622 | 6023 |
| CD8 naive | 5297 | 5337 | 5438 | 5363 | 5456 | 5419 | 5039 | 4959 | 5384 | 5499 | 5570 | 5526 | 5240 | 5021 | 5322 | 5127 | 5068 | 5287 |
| CD8 TCM | 7431 | 7804 | 8297 | 7789 | 8408 | 8565 | 5578 | 4975 | 5863 | 7632 | 5526 | 9052 | 5590 | 5510 | 5754 | 5604 | 5802 | 6207 |
| CD8 TEM | 5341 | 5367 | 5457 | 5368 | 5465 | 5435 | 5040 | 4882 | 5460 | 5517 | 5240 | 5590 | 5606 | 5310 | 5374 | 5373 | 5297 | 5436 |
| CD8 TEMRA | 5274 | 5300 | 5377 | 5269 | 5368 | 5334 | 4902 | 4726 | 5257 | 5384 | 5021 | 5510 | 5310 | 5524 | 5183 | 5287 | 5361 | 5364 |
| Treg | 5511 | 5549 | 5659 | 5543 | 5658 | 5639 | 5141 | 4939 | 5609 | 5714 | 5322 | 5754 | 5374 | 5183 | 5780 | 5257 | 5252 | 5462 |
| MAIT | 5352 | 5378 | 5477 | 5371 | 5474 | 5447 | 4964 | 4812 | 5404 | 5495 | 5127 | 5604 | 5373 | 5287 | 5257 | 5627 | 5303 | 5424 |
| CD56 dim NK | 5531 | 5584 | 5659 | 5540 | 5664 | 5592 | 5035 | 4720 | 5312 | 5622 | 5068 | 5802 | 5297 | 5361 | 5252 | 5303 | 5829 | 5649 |
| CD56 bright NK | 5924 | 5983 | 6093 | 5967 | 6121 | 6028 | 5280 | 4833 | 5535 | 6023 | 5287 | 6207 | 5436 | 5364 | 5462 | 5424 | 5649 | 6265 |

**Table S4. Number of expressed autosomal genes shared between each designated pair of PBMC cell types, related to Figure 3.** Genes must have a non-zero count in at least 10% of the cells for the given PBMC cell type to be considered expressed.
